# The impact of wild-boar derived microbiota transplantation on piglet microbiota, metabolite profile and gut proinflammatory cytokine production differs from sow-derived microbiota

**DOI:** 10.1101/2024.11.14.623678

**Authors:** Rajibur Rahman, Janelle M. Fouhse, Tingting Ju, Yi Fan, Tulika Bhardwaj, Ryan K. Brook, Roman Nosach, John Harding, Benjamin P. Willing

**Author notes:** Corresponding author: Department of Agricultural, Food & Nutritional Science, Faculty of Agricultural, Life & Environmental Sciences, University of Alberta, 116 St. and 85 Ave. Edmonton, Alberta T6G 2R3, Canada.

## Abstract

Colonization of co-evolved, species-specific microbes in early life plays a crucial role in gastrointestinal development and immune function. This study hypothesized modern pig production practices have resulted in the loss of co-evolved species and critical symbiotic host-microbe interactions. To test this, we reintroduced microbes from wild boars (WB) into conventional piglets to explore their colonization dynamics and effects on gut microbial communities, metabolite profiles, and immune responses. At postnatal day (PND) 21, 48 piglets were assigned to four treatment groups: 1) WB-derived mixed microbial community (MMC), 2) sow-derived MMC, 3) a combination of WB and sow MMC (Mix), or 4) Control (PBS). Post-transplantation analyses at PND 48 revealed distinct microbial communities in WB-inoculated piglets compared to Controls, with trends toward differentiation from Sow but not Mix groups. WB-derived microbes were more successful in colonizing piglets, particularly in the Mix group, where they competed with sow-derived microbes. WB group cecal digesta enriched with *Lactobacillus helveticus*, *Lactobacillus mucosae*, and *Lactobacillus pontis*. Cecal metabolite analysis showed that WB piglets were enriched in histamine, acetyl-ornithine, ornithine, citrulline, and other metabolites, with higher histamine levels linked to *Lactobacillus* abundance. WB piglets exhibited lower cecal IL-1β and IL-6 levels compared to Controls and Sow groups, while the Mix group showed reduced IFN-γ, IL-2, and IL-6 compared to the Sow group. No differences in weight gain, fecal scores, or plasma cytokines were observed, indicating no adverse effects. These findings support that missing WB microbes effectively colonize domestic piglets and may positively impact metabolite production and immune responses.

**Importance:** This study addresses the growing concern over losing co-evolved, species-specific microbes in modern agricultural practices, particularly in pig production. The implementation of strict biosecurity measures and widespread antibiotic use in conventional farming systems may disrupt crucial host-microbe interactions that are essential for gastrointestinal development and immune function. Our research demonstrates that by reintroducing wild boar-derived microbes into domestic piglets, these co-evolved microbes can successfully colonize the gut, influence microbial community composition, and alter metabolite profiles and immune responses without causing adverse effects. These findings also suggest that these co-evolved microbes can fill an intestinal niche, positively impacting immune activation. This research lays the groundwork for future strategies to enhance livestock health and performance by restoring natural microbial populations that produce immune-modulating metabolites.

**Graphical abstract:** 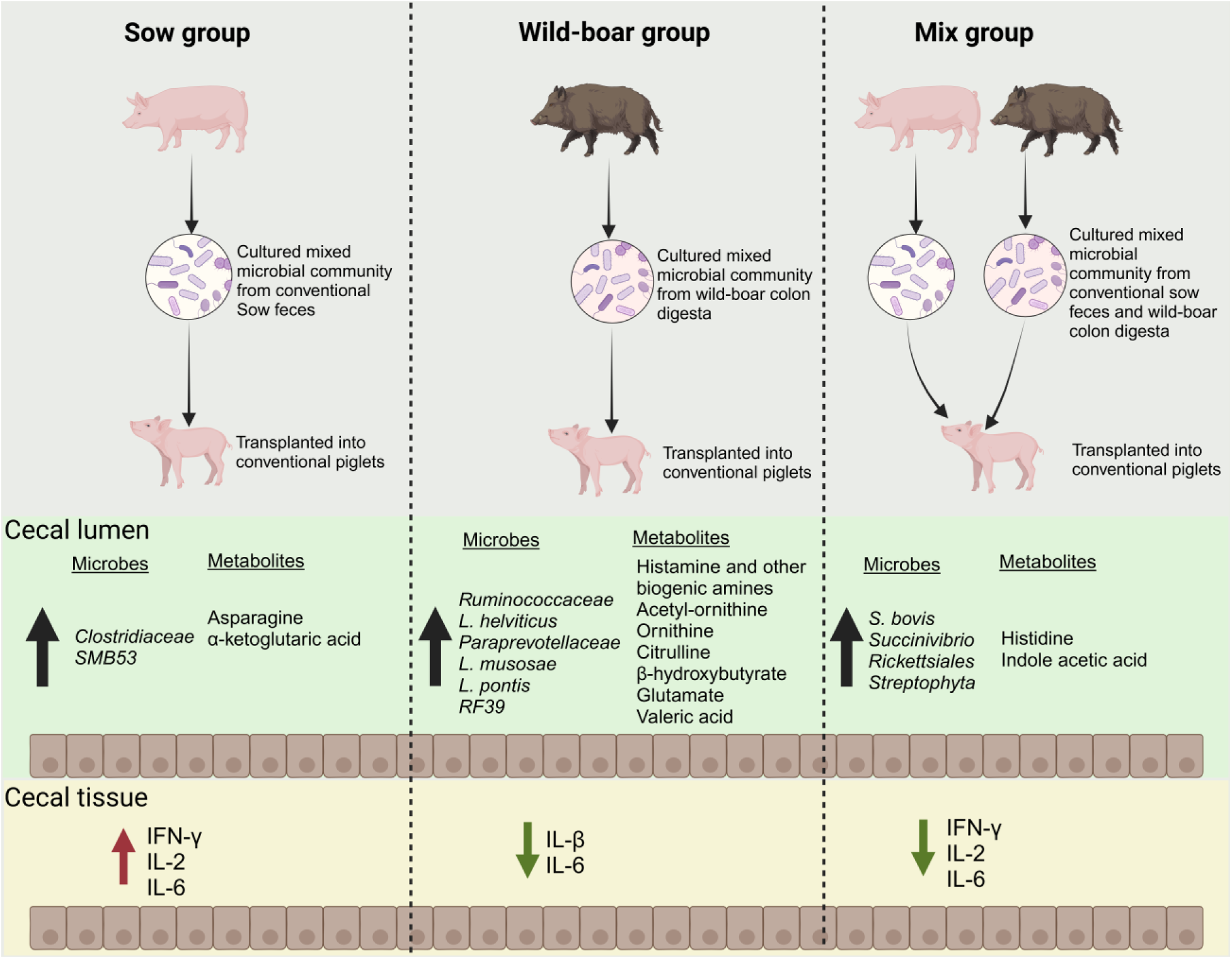

## Background

The mammalian gastrointestinal (GI) tract harbors a complex and dynamic community of bacteria, archaea, viruses, fungi, and other unicellular organisms, including protozoa and helminths, collectively known as the microbiome (1). These microbes colonize the gut through vertical transmission and environmental exposure, having co-evolved with their hosts to form a symbiotic relationship that is vital for host physiology (2). The combination of host and microbiome genes, referred to as the metagenome, largely shapes host physiology and biological processes.(3) Native commensal microbes support the host by producing beneficial metabolites, aiding energy metabolism, maintaining intestinal barrier function, protecting against pathogens, and regulating the immune system (4, 5). Importantly, early-life exposure to and colonization by these host-adapted microbes are essential for these beneficial outcomes (6–8).

In modern pig production, biosecurity measures such as the use of broad-spectrum disinfectants, pressure washing, and animal isolation help eliminate pathogens, but also inadvertently reduce pigs’ exposure to native commensal bacteria, leading to decreased gut microbial diversity (9, 10). Beyond sanitation practices, the microbiome of domestic pigs has evolved over relatively short periods within controlled environments, lacking exposure to diverse microorganisms (11, 12).

Numerous studies have reported that the gut microbial composition of wild animals, including pigs, differs from that of their domestic or laboratory counterparts (11, 13, 14). This lack of indigenous microbes can impair host intestinal and immune development, lowering resistance to pathogens (15–17).

During weaning, piglets often experience disruptions in their gut microbial community due to dietary and environmental changes, leading to dysbiosis and an increased risk of intestinal infections and related health issues (18, 19). Nearly 10% of piglet deaths during the weaning transition have been attributed to diarrhea, a complication associated with an underdeveloped gut microbiome and immune system, resulting in economic loss and welfare concerns in the pig industry (20). To address these challenges, antibiotics are frequently administered in piglets during weaning, both therapeutically and prophylactically. However, antibiotic use can worsen the situation by further disrupting the gut microbiome, weakening gut health, and contributing to the rise of antimicrobial resistance, a major global issue (21–23). Fecal Microbiota Transplantation (FMT) has emerged as a promising alternative for restoring microbial balance and intestinal health, potentially making piglets more resilient to external stressors. FMT involves transferring fecal matter containing viable microbes from a healthy donor to the recipient’s GI tract to restore dysbiosis and repopulate the gut with beneficial bacteria (24). FMT has successfully treated recurrent and refractory *Clostridioides difficile* infections in humans (25). However, its effectiveness in treating other conditions, such as inflammatory bowel disease, has been inconsistent (26, 27). Similarly, FMT outcomes in pigs have varied, with some studies reporting improvements in growth, intestinal development, and immune system maturation in weanling piglets (28–30), while others have observed reduced body weight and poor intestinal development (31, 32).

Several factors may contribute to the varying FMT outcomes in human and animal studies. Negative associations between donor fecal components, such as pathogenic bacteria, viruses, and fungi, can impact FMT efficacy (33–35). Microbial composition variations among donors and bacteria’s viability in the inoculum are also critical for FMT success (36, 37). A recent study from our group showed that using a cultured, mixed microbial community inoculum, rather than direct FMT, was more effective at modulating gut microbial composition, excluding pathogens, and reducing donor variability (8). The success of FMT is also linked to the colonization of specific bacterial strains and their metabolic capacities. Previous human FMT studies have highlighted the importance of colonizing specific bacterial strains with the metabolic capacity to produce beneficial metabolites for long-term success (38, 39). Another challenge is that the microbiota of conventional pigs, like laboratory mice, lacks resilience and can change with minor stressors, such as weaning, changing diets, or even moving to a different pen within the same facility (40–42). A recent study in mice reported that transferring wild mouse microbiota into lab mice improved microbiome stability and increased resilience to environmental stressors (15). Another study showed that wild mouse microbiota transplants could enhance laboratory mice’s infection resistance and disease resilience (16). Therefore, using a defined, naturally co-evolved microbial community instead of whole fecal matter may improve the consistency and efficacy of FMT by colonizing host-adapted beneficial bacteria.

This explorative FMT study aimed to investigate the interactions between microbes and their hosts following gut microbiota transplantation from wild boar, domestic pigs, and a combination of both into weaning piglets. The goals were to characterize microbial colonization, assess changes in gut microbiota composition, investigate metabolite profiles, and evaluate host immune responses. The study included (i) tracking temporal changes in the piglet gut microbiota post-transplantation using metataxonomic analysis, (ii) identifying gut metabolite alterations associated with transplantation using targeted metabolomics, and (iii) evaluating the host’s response to microbial transplant by measuring growth performance, fecal scores, plasma cytokines, lactate levels, and cecal cytokines.

## Methods and materials

### Ethics statement

This animal trial was designed and conducted following Canadian Council for Animal Care guidelines and approved by the University of Saskatchewan Animal Research Ethics Board (AUP20190075) and Animal Research: Reporting *In Vivo* Experiments (ARRIVE) guidelines. The study was conducted in biocontainment level 2 facilities at the University of Saskatchewan’s animal care unit, under the supervision of research staff trained through the mandatory University Animal Care Committee (UACC) ethics course in farm animals. The study was also approved by the University of Alberta’s Animal Care and Use Committee (AUP00002985). The capture and handling of wild pigs were conducted under protocols approved by the University of Saskatchewan (AUP20150024) and the Saskatchewan Ministry of Environment (permit 17FW027).

### Study design, Animal selection and housing

A total of 48 piglets from six different litters (n = 8/litter) were randomly selected at weaning on postnatal day (PND) 20 from Prairie Swine Centre (Saskatoon, SK). The piglets were transported to a biosafety level 2 facility at the Western College of Veterinary Medicine, University of Saskatchewan. Upon arrival, they were distributed across eight pens (n = 6/pen) with equal distribution of starting weight, sex, and litter of origin. Each room contained four pens (4 feet x 6 feet), with six piglets per pen and a 2-foot space between pens to prevent the spread of microbial communities. Piglets from each pen were randomly assigned to one of four experimental groups (n = 12 per group): (1) Control; (2) cultured mixed microbial community (MMC) from sow feces (Sow); (3) MMC from wild boar colon digesta (WB); and (4) MMC derived from both sow feces and wild boar colon digesta (Mix). The Control group received an inoculum of 1X phosphate-buffered saline (PBS) containing 20% glycerol and 0.05% L-cysteine. Each treatment group received 2 mL of their respective treatments on PND 21, 23, 25, and 27, administered orally using a syringe. Throughout the experimental period (total 28 days; PND 21 corresponded to experimental day 0, while PND 48 was experimental day 27), piglets had *ad libitum* access to unmedicated, antimicrobial-free starter feed (Co-op’s Whole Earth Pig Starter) and water. Room temperature was maintained at 22 ± 2.5°C with a 12-hour light/dark cycle. A detailed study timeline is shown in Figure 1A. Piglet weights were recorded on PND 21, 27, 34, 41, and 48. Diarrhea incidence and severity were assessed twice daily from PND 21 to 48 using a fecal consistency rubric as described by Barbosa et al (43), with fecal scores ranging from 0 to 4. Specifically, fecal consistency was categorized as follows: 0 = normal; 1 = soft, wet cement-like; 2 = watery; 3 = diarrhea with mucus; and 4 = diarrhea with blood. On PND 48, all piglets were euthanized to collect cecal and colon tissue and digesta, which were snap-frozen in liquid nitrogen and stored at -80°C for future analysis.

**Figure 1:**
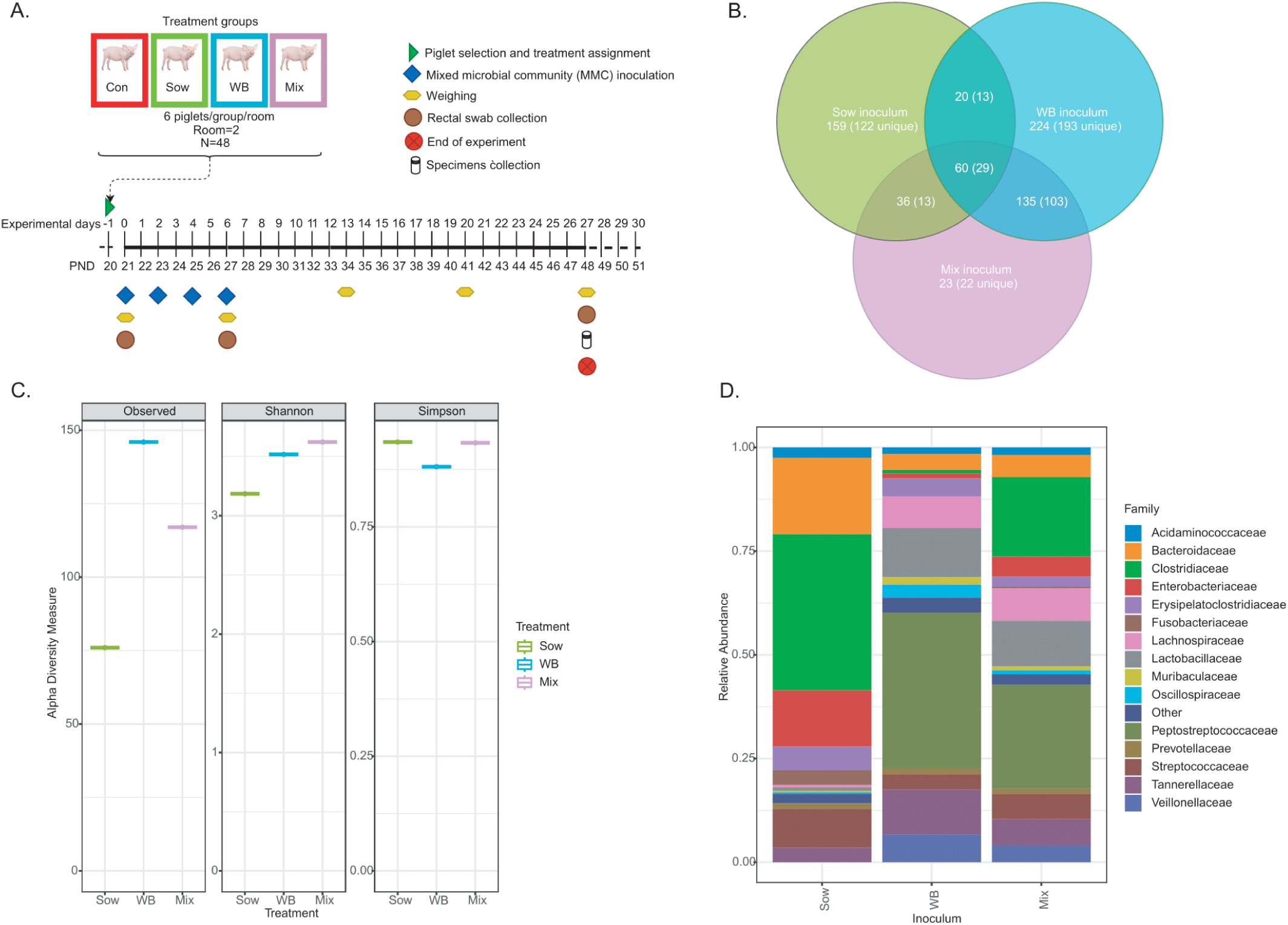
Experimental design and microbial profiling of inocula. (A) Schematic representation of experimental design and timeline. (B) α-diversity indices analysis of inocula showed that WB and Mix groups had higher microbial richness (Observed features) and diversity (Shannon) compared to Sow inoculum. WB inoculum had lower evenness (Simpson) compared to Sow and Mix inocula. (C) Venn diagram showing total, unique and shared ASVs among the inocula. (D) Taxa bar plot showing the top 15 bacterial families present in the inocula.

### Live fecal microbial inoculum preparation and administration

For the Sow treatment, a fresh fecal sample was collected by digital stimulation from a healthy second-parity sow with no history of antibiotic use in the previous three months, housed at the Swine Research and Technology Centre at the University of Alberta. The WB inoculum was obtained by aseptically collecting fresh distal colon digesta from an adult wild pig (ID: 10C) from the Moose Mountain area of Saskatchewan, Canada. Within two hours of collection, both the sow and wild boar samples were homogenized and diluted 1:20 in 1X PBS with 20% glycerol and 0.05% L-cysteine (w/v) and stored at -80°C. The samples were subsequently plated on four different media: fastidious anaerobe agar (Neogen, ref. NCM0014A), Difco™ brain heart infusion agar (BD, ref. 241820), Columbia blood agar base (Oxoid, ref. CM0331) with sheep blood (Thermo Fisher, ref. R54008), and Wilkins–Chalgren agar (Oxoid, ref. CM0642) in an anaerobic chamber with a gas mixture of 90% N2, 5% H2, and 5% CO2 (BACTRON 300; Sheldon Manufacturing, Cornelius, OR, USA). After 48 hours of incubation at 37°C, colonies from 10^−2^ and 10^−3^ dilution plates were pooled and suspended in 2 ml of 1X PBS with 20% glycerol and 0.05% L-cysteine. A portion of the pooled inoculum was plated anaerobically on fastidious anaerobe agar for bacterial enumeration, while the remaining was stored at -80°C until inoculation. Before administration, all inocula were diluted 1:1 with 1X PBS containing 20% glycerol and 0.05% L-cysteine (v/v) to achieve a final concentration of 1 × 10^8^ colony-forming units (CFUs)/mL. The mixed inoculum for the Mix group was prepared by combining the final Sow and WB inocula in a 1:1 ratio. All plating and dilution steps were performed in an anaerobic chamber with the same gas mixture.

### Sample collection

Fecal swabs were collected from all piglets prior to inoculation on PND 21 and post-inoculation on PND 27 and 48 using sterile cotton swabs, which were stored at -80°C until further analysis. Blood samples were collected on PND 48 from the external jugular vein into EDTA tubes pre- loaded with 200 µL of protease inhibitor. Plasma was separated by centrifugation (2,000 rpm, 4°C, 10 min), transferred into 1.5 mL microcentrifuge tubes, and snap-frozen in liquid nitrogen until further analysis. All pigs were humanely euthanized at PND 48 using a cranial captive bolt followed by exsanguination. Cecal (tip of the cecum) tissues and digesta were collected into sterile microcentrifuge tubes and immediately snap-frozen in liquid nitrogen for further analysis.

### Gut microbiota profiling

Total DNA was extracted from fecal swabs, cecal digesta, and inocula using the DNeasy PowerSoil kit (Qiagen®, Valencia, CA) following the manufacturers’ instructions with an additional bead-beating step described previously.(44) DNA concentrations were determined using the Quant-iT™ PicoGreen® dsDNA Assay Kit (Thermo Fisher Scientific, Waltham, MA, USA), and diluted to 5 ng/µl. Amplicon libraries targeting the V3–V4 region of the 16S rRNA gene were prepared using the Illumina 16S metagenomic sequencing library preparation protocol. The pooled library was then sequenced on an Illumina MiSeq platform (Illumina, Inc., San Diego, CA) with paired-end reads of 2 × 300 bp for 600 cycles, using the following PCR primers (341F/785R): Forward: 5′-TCGTCGGCAGCGTCAGATGTGTATAAGAGAC AGCCTACGGGNGGCWGCAG-3′; Reverse: 5′-GTCTCGTGGGCTCGGAGAT GTGTATAAGAGACAGGACTACHVGGGTATCTAATCC-3′. Sequencing data were processed using the Quantitative Insights Into Microbial Ecology 2 (QIIME2) pipeline (version 2024.2) (45). Quality control was performed using FastQC with default parameters, and sequences with an average quality score below 20 were trimmed. Forward and reverse sequences were truncated at 260 bp and 220 bp, respectively. The Divisive Amplicon Denoising Algorithm 2 (DADA2) plugin was employed for denoising and generating feature tables with default settings (46). Amplicon sequence variants (ASVs) were aligned using MAFFT (47), and a phylogenetic tree was constructed with FastTree2. Taxonomy assignment for ASVs was performed using the q2-feature-classifier (48) with the classify-sklearn Naive Bayes taxonomy classifier (49), trained on nearly complete 16S rRNA sequences from the SILVA database version 138 (50).

### Cecal metabolite profiling

A total of 70 metabolites (Supplementary Table 1) were profiled from 48 cecal digesta samples at The Metabolomics Innovation Center (TMIC) using the PRIME Assay kit. Samples were homogenized with extraction buffer A (85% methanol, 15% phosphate buffer) for lipids, amino acids, biogenic amines, glucose, and acylcarnitines, or with extraction buffer B (15% methanol, 85% phosphate buffer) for organic acids. After centrifugation, supernatants were collected for further analysis.

For most metabolites, 20 µl of internal standard (ISTD) and 10 µl of supernatant were dried, derivatized with 5% phenyl isothiocyanate, and reconstituted for liquid chromatography with tandem mass spectrometry (LC-MS/MS) and direct flow injection-mass spectrometry (DFI-MS/MS) analysis. For lipids, 5 µl each of ISTD and sample were mixed with 490 µl of DFI buffer and analyzed. Organic acids were extract using 10 µl of sample supernatant and 40 µl of isotope-labeled ISTD, derivatized with 3-nitrophenylhydrazine, stabilized, and diluted for LC-MS analysis. Metabolite profiling was performed using an Agilent 1260 ultra-high-pressure liquid chromatography (UHPLC) system (Agilent Technologies, Palo Alto, CA, USA) and AB SCIEX QTRAP® 4000 mass spectrometer (Sciex Canada, Concord, ON, Canada).

### Plasma and cecal cytokine analysis by a Porcine 13-plex cytokine/chemokine array

A 13-plex panel of porcine cytokines/chemokines was analyzed from plasma and cecal protein samples by Eve Technologies (Calgary, AB). Protein concentration in each sample was quantified using the Pierce™ BCA Protein Assay Kit (Thermo Fisher Scientific) and normalized to 3 mg/ml. The panel included Granulocyte–Macrophage Colony-Stimulating Factor (GM-CSF), Interferon-γ (IFN-γ), Tumor Necrosis Factor-alpha (TNF-α), and Interleukins (IL)-1α, IL- 1β, IL-1ra, IL-2, IL-4, IL-6, IL-8, IL-10, IL-12, and IL-18. Briefly, uniquely color-coded polystyrene beads were conjugated with capture antibodies for each cytokine/chemokine. Beads were incubated with samples in a 96-well plate, and the multiplex immunoassay was analyzed using a BioPlex 200 system, which measured fluorescent intensity signals to quantify cytokine concentrations. The results were calculated using a standard curve and expressed in pg/mL.

### Plasma Lactate quantification

Plasma D-Lactate levels were quantified using a D-Lactate Assay Kit (Sigma, ref: MAK336) following the manufacturer instructions. Briefly, 20 µl of both standards and plasma samples were transferred into designated wells of a 96-well clear flat-bottom plate. Subsequently, 80 µl of the reaction mix was quickly added to each well, and the plate was mixed thoroughly. Initial absorbance was measured immediately at 565 nm, followed by a 20-minute incubation at room temperature. After incubation, final absorbance was measured at 565 nm.

### Data visualization and statistical analysis

All statistical analyses were performed using GraphPad Prism (v9.5.1) and R (v3.5.2), and data were presented as mean ± standard error of the mean (SEM). A linear mixed-effect model was applied to compare body weight, fecal scores, and cytokine/chemokine levels among treatments, with sow and pen used as random effects. The presence of ASVs was visualized using ggplot2 package in R (v4.0.5). To evaluate successful bacterial transfer from the MMC inocula to recipients, ASVs from the inocula, piglets, or both were quantified and visualized. To reduce false positives, only ASVs with counts greater than two were included in the analysis. Changes in microbial community structure and α -diversity were visualized using the R package phyloseq. Alpha diversity indices, including Observed, Simpson, and Shannon indices, were measured and analyzed with significance by Kruskal-Wallis and post-hoc Dunn tests. Rarefication was performed based on the sample with the lowest sequence depth. Beta-diversity was analyzed based on Bray-Curtis dissimilarity and visualized using principal-coordinate analysis (PCoA).

Differences between microbial communities were assessed using the adonis function in the vegan package in R.(51) The differential abundance of key taxa at the genus level among treatment groups was analyzed using Linear discriminant analysis (LDA) effect size (LEfSe) package in R. Statistical significance was set at *P* ≤ 0.05, with 0.05 < *P* ≤ 0.10 considered indicative of a trend. Cecal metabolite analyses were performed using MetaboAnalyst 6.0 (https://www.metaboanalyst.ca/). Spearman correlation was used to assess relationships between cecal microbes and metabolites. Short-chain fatty acids, biogenic amines, polyamines, and cytokine data were log-transformed and analyzed with a linear mixed-effects model, accounting for the pen and litter effect as random variables. Procrustes analysis was performed on the Euclidian distances of eigenvalues for both the microbiome and metabolome datasets using the *Protest* function in the *vegan* R package.

## Results

### WB inoculum is composed of diverse and beneficial bacterial families

The WB inoculum exhibited a numerically higher bacterial diversity and richness (based on Observed and Shannon indices) than the Sow inoculum, although it had lower evenness based on Simpson indices (Figure 1B). The Mix inoculum had higher Shannon diversity than the WB and Sow inocula and showed higher Simpson evenness than the WB inoculum. The WB and Sow inocula contained 439 and 275 ASVs, respectively (Figure 1C). The inocula ASVs were compared to those found in the piglets and considered to be common: 101 in WB and 98 in Sow. Inocula ASVs that were not present in all piglets before inoculation were defined as “unique”, and further described as specific to the respective sources of inocula (i.e. WB or Sow). WB inoculum had higher unique ASVs than SOW, 338 vs 177 (Fisher’s exact test *P* = 0.0003). Of the unique ASVs in the inocula, 42 were shared between WB and SOW. At the family level, the WB inoculum was enriched with *Peptostreptococcaceae*, *Lactobacillaceae*, *Tannerellaceae*, *Lachnospiraceae*, *Veillonellaceae*, and *Erysipelatoclostridiaceae* (Figure 1D), whereas the Sow inoculum was enriched with *Clostridiaceae*, *Bacteroidaceae*, *Enterobacteriaceae*, *Streptococcaceae*, *Erysipelatoclostridiaceae*, and *Veillonellaceae*.

### FMT modulates microbial community structure and shapes gut microbiota composition toward carbohydrate degradation

Pre-transplantation (PND 21) fecal microbiota analysis revealed no differences in β-diversity based on Bray-Curtis dissimilarity matrix (Adonis *P* = 0.73, R^2^ = 0.05, beta-dispersion *P* = 0.22) or α -diversity indices based on Observed (*P* = 0.67), Shannon (*P* = 0.21) and Simpson indices (*P* = 0.18) (Figure 2A-B). By day 6 post-transplantation, there was a trend in β-diversity difference (Adonis *P* = 0.07, R^2^ = 0.07, beta-dispersion *P* = 0.90) among the treatments (Supplementary Figure 1A). WB inoculum recipients showed a trend of greater distance in microbial community structure to Sow (*P* = 0.09) and Control (*P* = 0.09) piglets than to the Mix group (*P* = 0.73). No differences were observed (*P* > 0.1) when comparing Control vs Sow, Control vs Mix, and Sow vs Mix groups. Alpha-diversity indices remained unchanged at day 6 post-transplantation (Supplementary Figure 1B).

**Figure 2.**
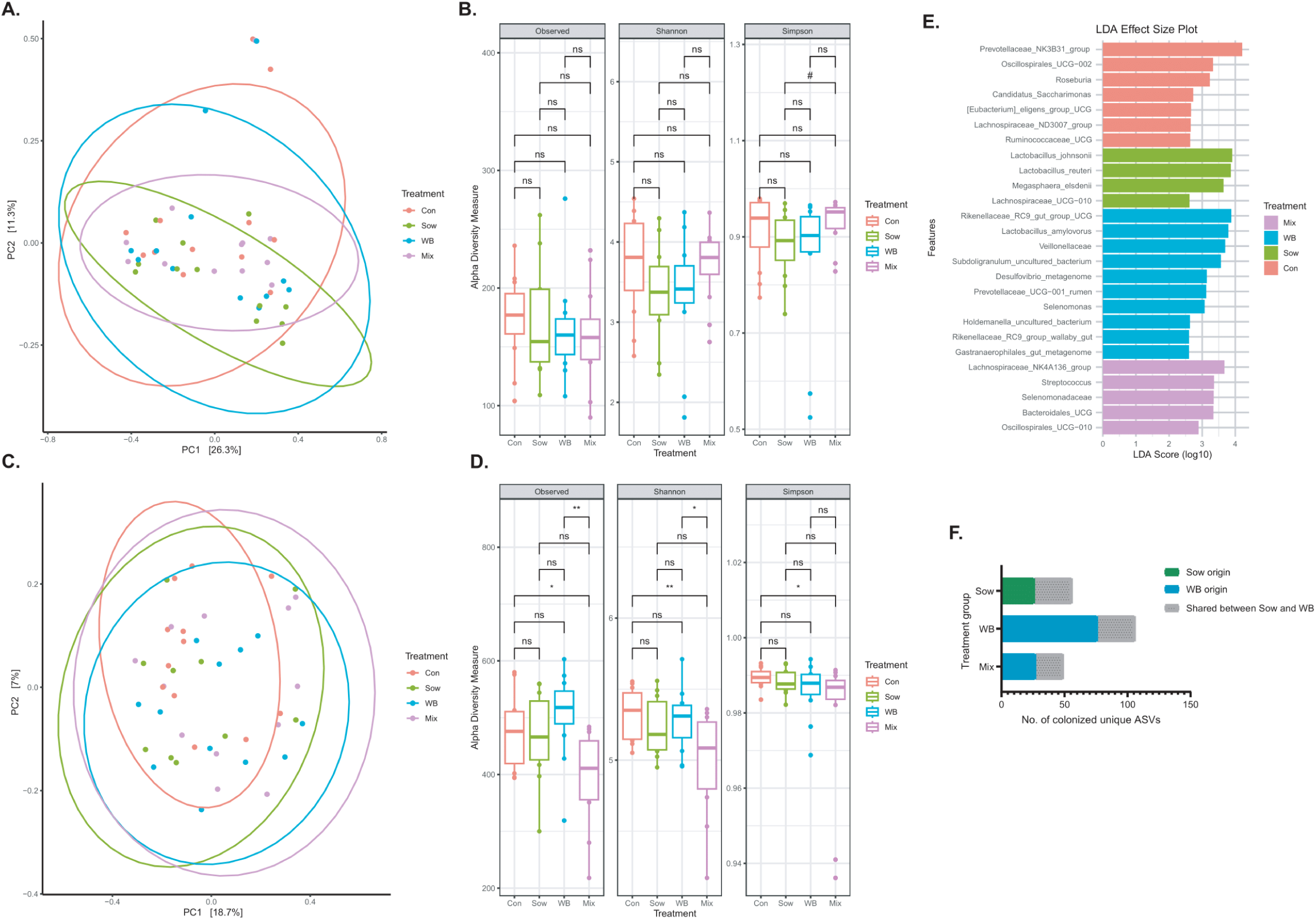
Comparison of fecal microbiota composition and α-diversity among piglets pre- transplantation and 28 days post-transplantation (A) Pre-transplantation fecal microbial community structure did not differ among groups (Sow, WB, Mix, and Control; n = 12/group), as measured by β-diversity using Bray–Curtis dissimilarity (Adonis, R² = 0.05, *P* = 0.73; Betadispersion *P* = 0.22). (B) No differences in species richness, diversity, or evenness were observed among groups before transplantation (KW test, Observed *P* = 0.67; Shannon *P* = 0.21; Simpson *P* = 0.18). (C) On day 28 post-transplantation (PND 48) post-transplantation, the overall fecal microbial community structure was altered (Sow: n = 10; WB: n = 12; Mix: n = 12; and Control: n = 12), with differences detected by β-diversity analysis (Adonis, R² = 0.08, *P* = 0.04; Betadispersion *P* = 0.50). (D) On day 28 post-transplantation (PND 48) α--diversity analysis indicated that the Mix group had lower species richness (Observed) and diversity (Shannon) than Control and WB groups (KW test, Observed *P* = 0.009; Shannon *P* = 0.04; Simpson *P* = 0.20). (E) On day 28 post-transplantation (PND 48), LEfSe analysis identified differentially abundant taxonomical features among treatment groups (*P* < 0.05). (F) Comparison of the presence of inoculum-originated unique ASVs among the respective recipient piglets at post-transplantation day 28 revealed WB-origin ASVs were more successful at engrafting (Fisher’s exact test *P* = 0.03). \**P* < 0.05, \*\**P* < 0.01, \*\*\**P* < 0.001, *^#^P* > 0.05 and < 0.1

LefSe revealed bacteria enrichment patterns on day 6 post-transplantation. WB inoculum recipients were enriched with a group of bacteria associated with fiber degradation and SCFA production, including *Prevotella*, *Treponema porcinum*, *Prevotellaceae*, *Selenomonas*, *WPS-2*, *Elusimicrobium*, *Bifidobacterium adolescentis*, *Bifidobacterium thermophilum*, *Veillonella caviae*, and *Selenomonadaceae* (Supplementary figure 1C). Sow inoculum recipients had increased levels of carbohydrate-fermenting bacteria including *Frisingicoccus*, *Sutterella*, *Lachnospiraceae UCG-003*, *[Eubacterium] coprostanoligenes* group, uncultured *Ruminococcaceae*, *Solobacterium*, and *Mailhella*. Interestingly, Control piglets displayed a higher abundance of potential pathogens such as *Campylobacter*, and carbohydrate-fermenting taxa such as *Desulfovibrio*, *Erysipelotrichaceae*, *[Eubacterium] hallii group*, and *Alloprevotella* (FDR *P* < 0.05). The Mix group showed similar enrichment patterns to the WB and Sow groups, with elevated levels of *Clostridia*, *[Eubacterium]_xylanophilum_group*, and *RF39*.

By post-transplantation day 28 (PND 48), there was a shift observed in microbial community structure (Adonis *P* = 0.04, R^2^ = 0.08, beta-dispersion *P* = 0.50), with the WB group showing distinct differences from the Control group (*P* = 0.02) and a trend toward divergence from the Sow group (*P* = 0.07), but not from the Mix group (*P* = 0.23) (Figure 2C). Moreover, the Mix group showed a trend toward difference from the Control group (*P* = 0.08), but not the Sow group (*P* = 0.22). The α-diversity indices on post-transplantation day 28 (PND 48) revealed that Observed features (*P* = 0.009) and Shannon index (*P* = 0.04) were different among treatment groups (Figure 2D). The WB and Control groups had higher Observed and Shannon diversity than the Mix group.

LEfSe analysis at PND 48 revealed that WB piglets were enriched with taxa including *Rikenellaceae RC9 gut group*, *Lactobacillus amylovorus*, *Veillonellaceae*, *Subdoligranulum UCG*, *Desulfovibrio*, *Prevotellaceae UCG-001*, *Selenomonas*, *Holdemanella UCG*, *Rikenellaceae RC9 gut group*, and *Gastranaerophilales* (Figure 2E). The Sow group was enriched with *Lactobacillus johnsonii*, *Lactobacillus reuteri*, *Megasphaera elsdenii*, and *Lachnospiraceae UCG-010*. The Control piglets showed enrichment in *Prevotellaceae NK3B3*, *Oscillospirales UCG-002*, *Roseburia*, *Candidatus Saccharimonas*, *[Eubacterium] eligens group*, *Lachnospiraceae ND3007 group*, and *Ruminococcaceae UCG*. The Mix group was enriched with *Lachnospiraceae NK4A136*, *Streptococcus*, *Selenomonadaceae*, *Bacteroidales UCG*, and *Oscillospirales UCG-01*.

### WB inoculum-derived unique ASVs were more successful in colonizing the piglet gut

The successful colonization of unique ASVs was evaluated (Figure 2F). At day 28 post-transplantation, 106 unique ASVs derived from the WB inoculum were detected in the WB group, with a relative abundance ranging from 1.4% to 18.9% of the total community (mean: 6.5%). Of these, 30 ASVs were shared with Sow inoculum (relative abundance range: 0.2% to 13%; mean: 3.7%). 56 unique ASVs derived from the Sow inoculum were detected in the Sow piglets, with a relative abundance range of 2% to 6.6% (mean: 4.5%). Of the 30 ASVs that were shared with the WB inoculum relative abundance ranged from 0.2% to 5.4% in the sow group (mean: 2.9%). 49 unique ASVs in the WB inoculum were detected in the Mix group (relative abundance range: 0.3% to 12.1%; mean: 4.0%) of the community), 22 of which were shared between WB and Sow (relative abundance range: 0.07% to 10.4%; mean: 2.3%), whereas no unique ASVs specific to Sow were detected (Fisher exact test *P* = 0.03). The results identify successful colonization of ASVs from the inocula, with greater success of those from WB or shared between WB and Sow.

### FMT did not alter growth performance, fecal scores and plasma lactate and cytokines among inocula recipients

FMT did not influence the growth performance of piglets based on mean body weight gain (Figure 3A; *P* = 0.55). Additionally, fecal consistency scores remained consistent across all groups during the experiment (Figure 3B; *P* = 0.87). No differences were found in plasma lactate levels among groups measured on PND 48 (Figure 3C; *P* = 0.81). There were no differences in plasma cytokines and chemokines among treatment groups (Figure 3D; *P > 0.05*).

**Figure 3:**
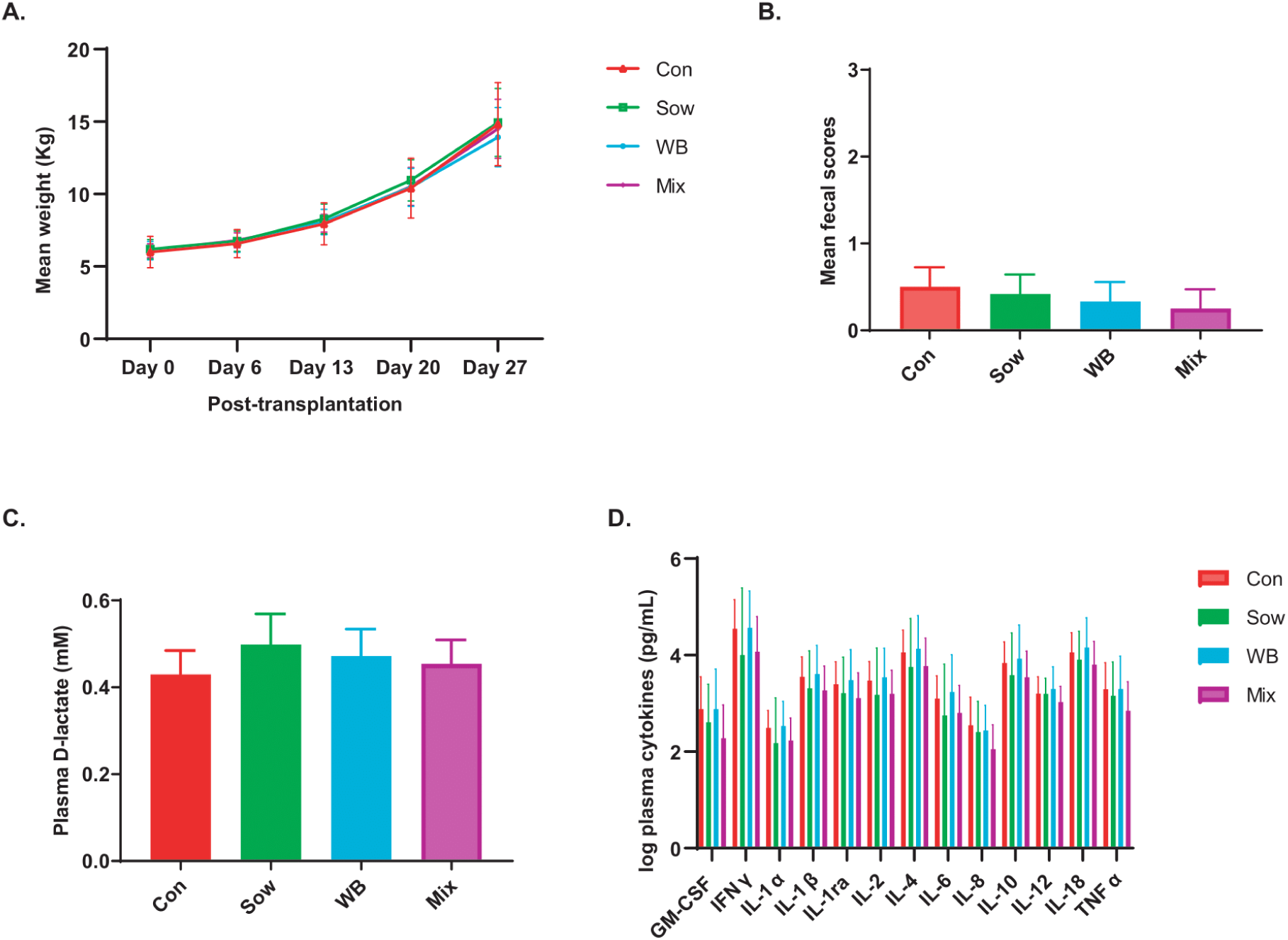
The effects of FMT on piglet weight gain, fecal score as well as plasma D-Lactate and cytokine/chemokine levels. Comparison of (A) weight gain (n = 12/group), (B) fecal score (n = 12/group), (C) Plasma D- Lactate (n = 5/group), and (D) cytokines/chemokines indicated no differences among treatment groups (n = 12/group). All data are shown as Mean with SEM. Statistical analysis was conducted to examine the treatment effect using linear mixed effect models (where litter and pen effect accounted for a random effect) followed by pairwise comparison. \**P* < 0.05, \*\**P* < 0.01, \*\*\**P* < 0.001, *^#^P* > 0.05 and < 0.1

### Wild boar-derived FMT inocula recipient piglets showed an altered cecal microbial community structure and composition

At PND 48 (28 days post-transplantation), a shift in the cecal microbial community structure was observed (Adonis *P* = 0.02, R² = 0.09, beta-dispersion *P* = 0.23) among treatment groups (Figure 4A). Particularly, the Mix group exhibited a unique microbial community structure compared to the Sow group (*P* = 0.03) and showed a trend when compared to Control (*P* = 0.07) and WB (*P* = 0.08) groups. Interestingly, the Sow group did not differ from the Control group (*P* = 0.37). The WB group also showed a trend towards a different microbial community structure compared to both the Sow (*P* = 0.06) and Control groups (*P* = 0.09). No differences in Shannon (*P* = 0.25) and Simpson indices (*P* = 0.59) were observed among treatment groups (Figure 4B). However, the Mix group exhibited lower Observed features compared to the Control (*P* = 0.01) and WB (*P* = 0.02) groups.

**Figure 4.**
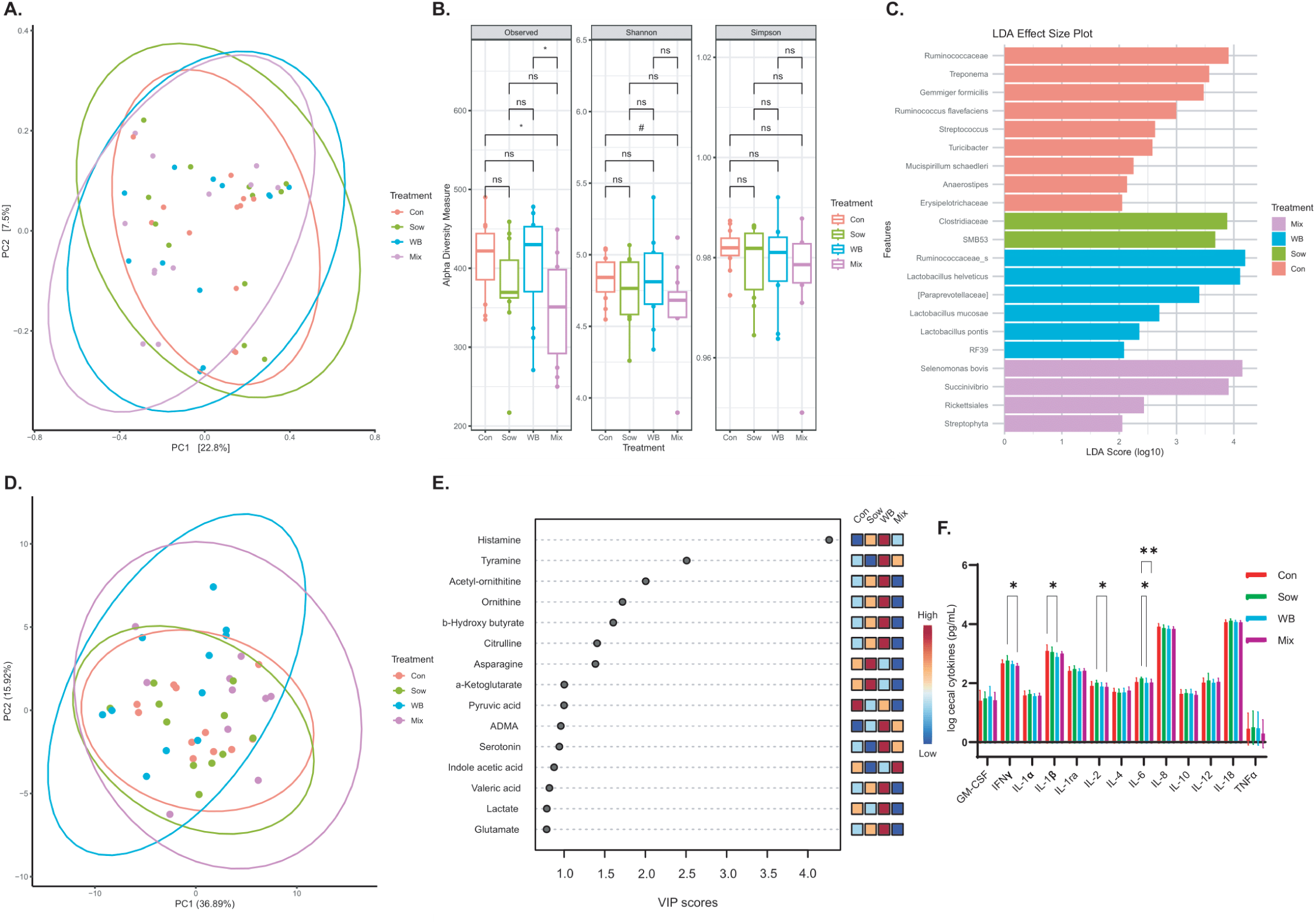
Comparison of post-transplantation day 28 (PND 48) cecal microbiota composition, α-diversity, and metabolite composition among treatment groups. (A) At day 6 (PND 27) post-transplantation, there was a shift in microbial community structure as measured by β-diversity based on Bray–Curtis dissimilarity (Adonis, R² = 0.09, *P* = 0.02; Betadispersion *P* = 0.23) (n = 12/group). (B) No differences were observed in Shannon or Simpson indices among treatment groups, however, the Mix group exhibited lower species richness (Observed features) compared to the Control (*P* = 0.01) and WB groups (*P* = 0.02) (Observed *P* = 0.03; Shannon *P* = 0.59; Simpson *P* = 0.38) (C) LEfSe analysis identified differentially abundant taxonomical features among treatment groups (*P* < 0.05). (D) Multivariate analysis of cecal metabolites at PND 48 revealed distinct clustering among treatment groups (Adonis *P* = 0.02, R² = 0.14; n=11/group). (E) VIP plot showing the top 15 metabolites enrichments across groups (n = 11/group). (F) Cecal cytokine/chemokine responses across groups (n = 12/group). Missing values were replaced with half the limit of detection (LOD/2). The data were log-transformed and analyzed using a linear mixed effects model, with the litter and pen effect included as a random factor. P-values were derived from adjusted pairwise comparisons, with significance set at P < 0.05. \**P* < 0.05, \*\**P* < 0.01, \*\*\**P* < 0.001, *^#^P* > 0.05 and < 0.1

LEfSe analysis of cecal digesta revealed distinct bacterial taxa enrichment patterns across treatment groups (Figure 4C). In the WB group, *Ruminococcaceae*, *Lactobacillus helveticus*, *[Paraprevotellaceae]*, *Lactobacillus mucosae*, *Lactobacillus pontis*, and *RF39* were enriched. The Sow group showed differential abundances of *Clostridiaceae* and *SMB53*. The Control group was enriched with *Ruminococcaceae*, *Treponema*, *Gemmiger formicilis*, *Ruminococcus flavefaciens*, *Streptococcus*, *Turicibacter*, *Mucispirillum schaedleri*, *Anaerostipes*, and *Erysipelotrichaceae*. The Mix group exhibited enrichment with *Selenomonas bovis*, *Succinivibrio*, *Rickettsiales*, and *Streptophyta*.

### WB piglets showed an altered cecal metabolite profile with enrichment of Lactobacillus-associated metabolites with lower pro-inflammatory cytokines

To investigate the metabolic shift linked to microbial colonization, targeted metabolomic profiling was conducted on cecal digesta samples. A multivariate analysis revealed distinct clustering (Adonis *P* = 0.02, R² = 0.14) among treatment groups (Figure 4D), with the WB group exhibiting differences from the Mix (*P* = 0.01) and Sow (*P* = 0.03) groups, and a trend compared to the Control group (*P* = 0.08). Interestingly, no difference was observed among the other groups. VIP analysis identified distinct metabolic enrichments across the groups (Figure 4E).

The WB group was notably enriched with histamine, acetyl-ornithine, tyramine, ornithine, b-hydroxy butyrate, citrulline, lactate, and glutamate, while the Sow group showed higher levels of asparagine and alpha-ketoglutarate. The Control group was enriched with pyruvic acid, and the Mix group had higher levels of creatine and indole acetic acid. Given that the WB group exhibited higher levels of histamine and tyramine, a subsequent analysis of other biogenic amines and histidine in the cecal digesta was performed (Figure 5A-H). Notably, the WB group showed elevated spermine levels and a trend towards higher serotonin and dopamine compared to the Sow group. Additionally, the Mix group displayed a trend of lower dopamine relative to the WB group. When examining histamine precursors, the WB group had lower histidine levels than the other three groups. The analysis of cytokines/chemokines in cecal tissue revealed that the Mix group had lower IFN-γ, IL-2, and IL-6 levels than the Sow group (Figure 4F). In contrast, the WB group exhibited a lower level of IL-1β compared to the Control group and a lower level of IL-6 compared to the Sow group.

**Figure 5:**
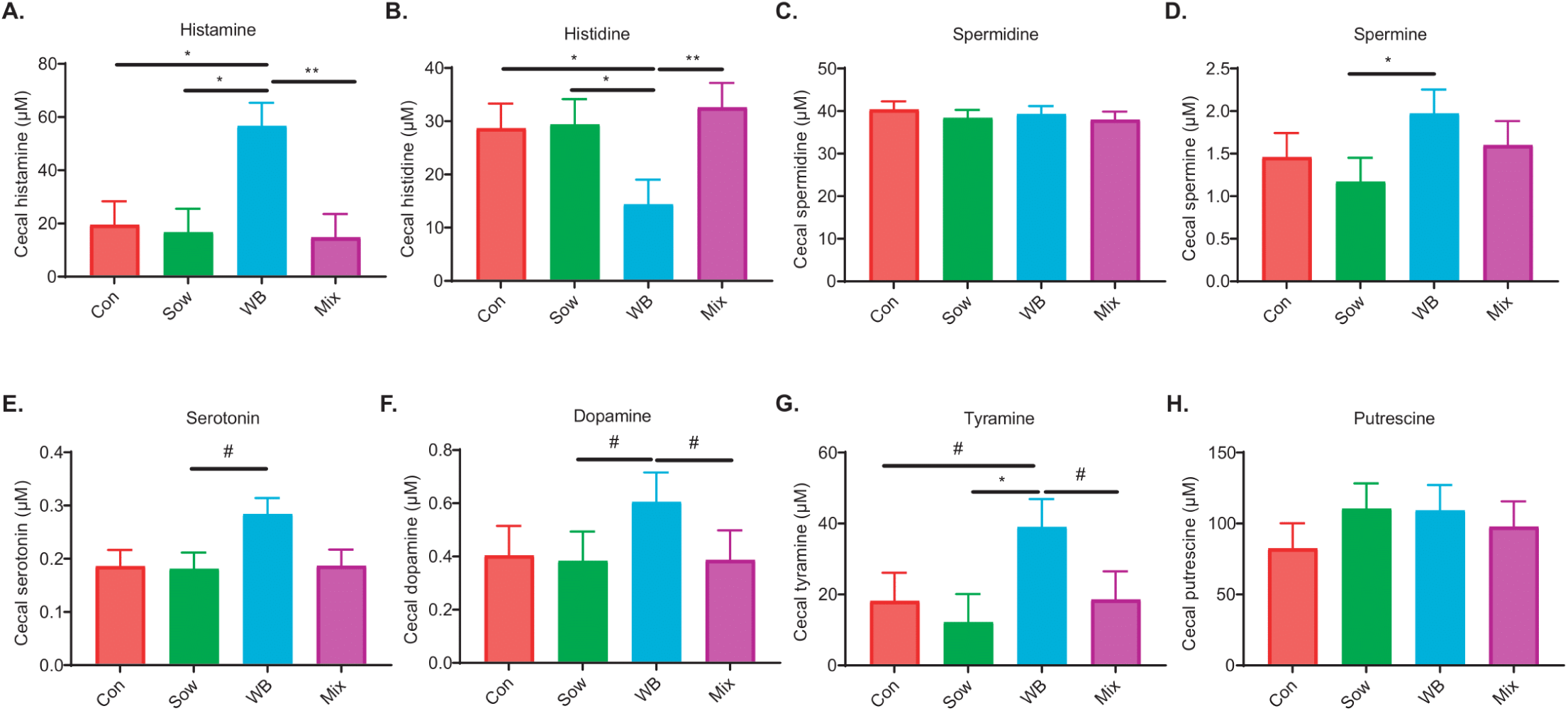
Comparison of cecal biogenic amines and histidine among treatment groups. (A) Histamine; (B) Histidine; (C) Spermidine; (D) Spermine; (E) Serotonin; (F) Dopamine; (G) Tyramine; and (H) Putrescine levels (n = 12/group). All data is shown as a mean with SEM. *P*- values were obtained from adjusted pairwise comparisons after a linear mixed effect model (where litter and pen effect accounted for a random effect), and significance was considered at *P <* 0.05. \**P* < 0.05, \*\**P* < 0.01, \*\*\**P* < 0.001, *^#^P* > 0.05 and < 0.1

No significant differences in butyrate, acetate, or propionate levels were observed among treatments. However, the WB group showed higher concentrations of valeric acid compared to the Mix group (Supplementary Figure 2A-J). There was no difference in branched-chain amino acids or intermediate energy metabolites such as pyruvate, succinate, and lactate.

Correlation analysis between cecal microbes and metabolites revealed that elevated levels of biogenic amines and metabolites, including histamine, tyramine, ornithine, and acetyl-ornithine, were positively associated with the abundance of cecal *Lactobacillus* (Figure 6). Interestingly, *Lactobacillus* abundance was negatively correlated with histidine levels in the cecum. Additionally, the genus SMB53 from the family *Clostridiaceae* was negatively associated with histamine levels. The association between the cecal microbiome and metabolome was further confirmed by applying Procrustes analysis on the PCoA axes of the two datasets (Procrustes sum of squares, 0.75; correlation, 0.49; P < 0.001) (Supplementary Figure 3).

**Figure 6:**
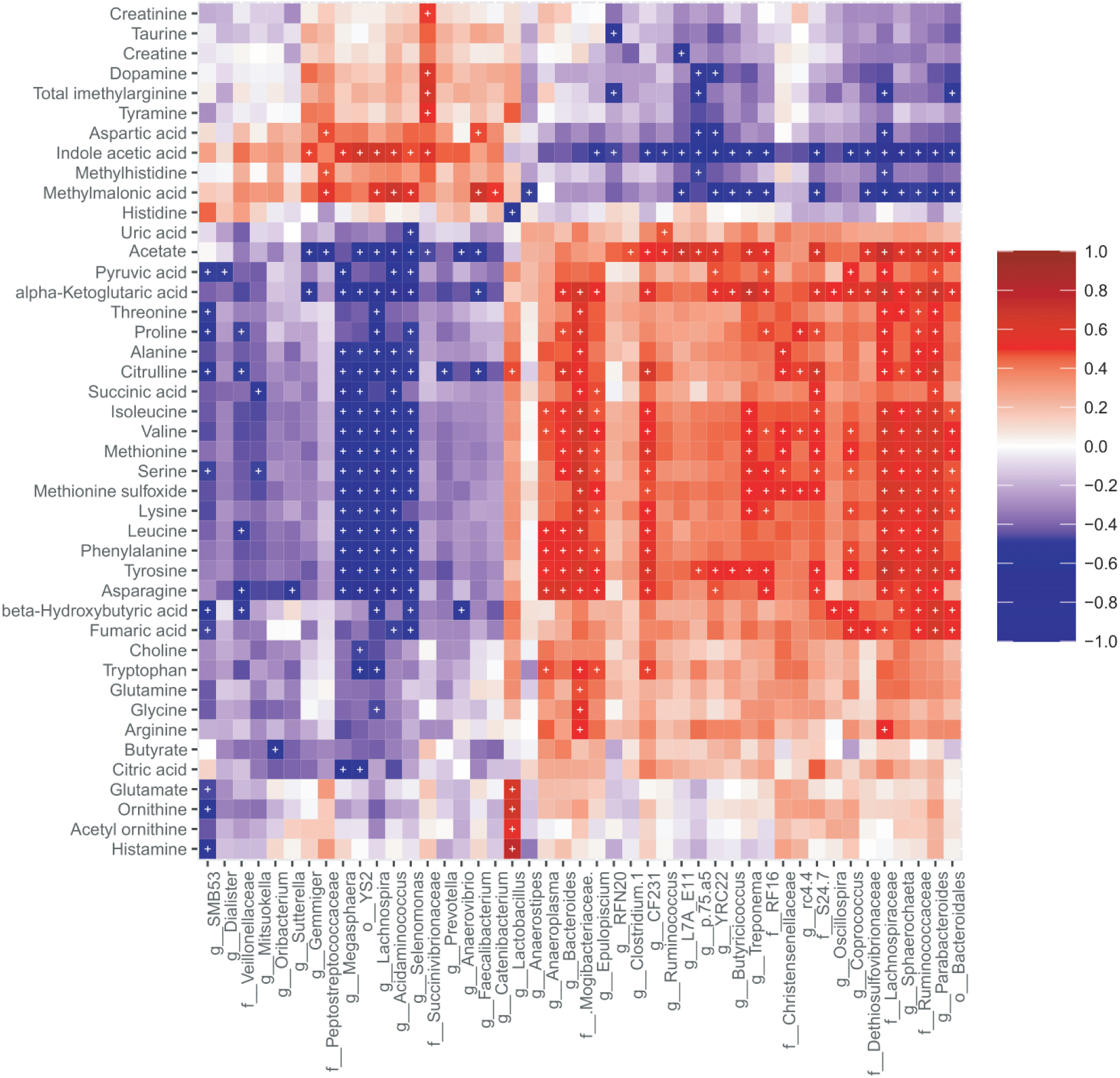
H**e**atmap **showing correlations between cecal microbes and metabolites.** Increased cecal histamine and other biogenic amines are associated with increased *Lactobacillus* abundance. Spearman’s correlation coefficients were used to calculate pairwise comparisons between microbial taxa abundances and metabolite concentrations. Microbe and metabolite found correlated after FDR adjustment <0.01 were marked with an asterisk.

### Metabolic pathway enrichment mirrored the cecal metabolites profile of WB groups

The Kyoto Encyclopedia of Genes and Genomes (KEGG) pathway enrichment analysis was used to explore the biological and metabolic functions of altered metabolites among inocula recipients (Figure 7). Compared to the Control group, Sow inoculum recipients were enriched in pathways related to glutathione metabolism, glycine, serine, and threonine metabolism, and the biosynthesis of neomycin, kanamycin, and gentamicin. In contrast, WB inoculum recipients showed enrichment in pathways associated with arginine biosynthesis, butanoate metabolism, histidine metabolism, arginine and proline metabolism, nitrogen metabolism, taurine and hypotaurine metabolism, and biotin metabolism. The Mix inoculum recipients were enriched in taurine and hypotaurine metabolism, as well as the degradation of valine, leucine, and isoleucine. All transplantation recipient groups demonstrated consistent enrichment in primary bile acid synthesis.

**Figure 7.**
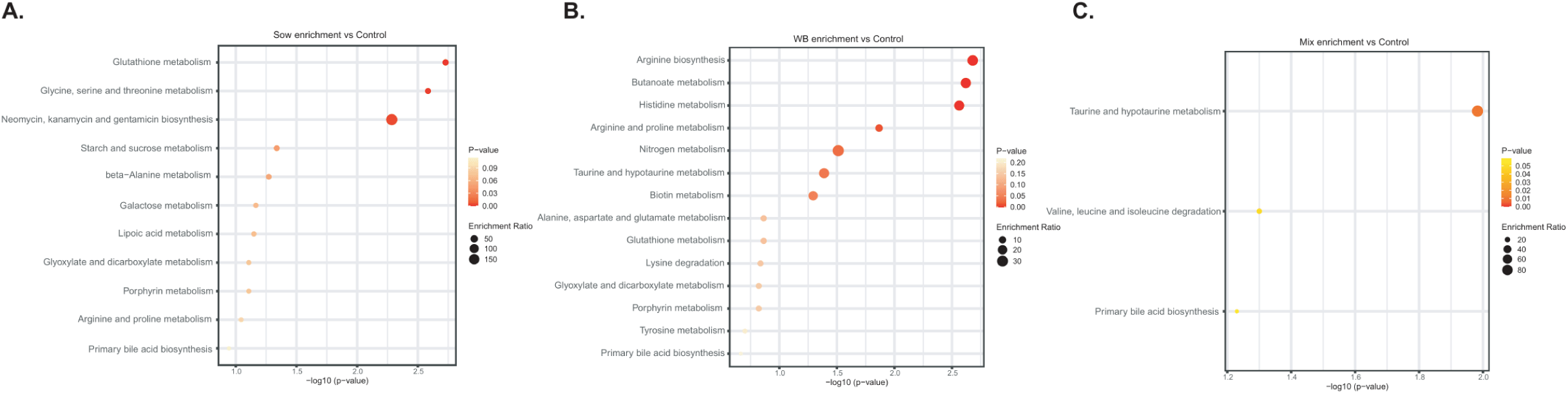
KEGG pathway enrichment analysis based on differentially abundant metabolites (*P* < 0.05) in the inocula recipients in PND 48 cecal digesta. (A) Control vs. Sow; (B) Control vs. WB; (C) Control vs. Mix.

## Discussion

Mammalian gut microbial communities have co-evolved with their hosts, playing essential roles in host physiology. These host-adapted microbes are critical for helping the host adapt to environmental changes and maintain resilience against pathogens (16, 52). Numerous studies have reported that microbial communities in wild animals are distinct from those in domesticated animals (14, 53, 54). Studies have also demonstrated that the gut microbiome of wild animals is associated with enhanced immune development and greater disease resilience compared to their domesticated counterparts (11, 28, 55, 56). In this study, cultured gut bacterial communities from wild and domestic pigs were transplanted into conventional post-weaning piglets, leading to significant changes in gut microbial composition, as well as cecal metabolite and cytokine profiles. Although FMT resulted in notable changes in gut microbiota and metabolite composition, no significant differences among the treatment groups were observed in weight gain, systemic cytokine profile, or fecal scores.

It has been documented that co-evolved microbes are better adapted to their hosts, giving them an advantage in colonizing the host gut compared to non-native microbes (57, 58). Our study found that the WB group’s microbial community structure began to diverge from the Control and Sow groups a week after inoculation and became distinct by day 28 post-transplantation (PND 48). Interestingly, the microbial community in the Control group remained similar to that of the Sow group, yet it differed from both the WB and Mix groups. This shift in the microbial community of the Sow group towards that of the Control group suggested a possible overlap in their compositions. On the other hand, despite receiving microbiota from sows, the Mix group’s divergence from the Control group may indicate the successful colonization of wild boar-derived microbiota. Similarly, a recent study in mice found that transplanting native microbes from wild mice resulted in altered microbial compositions compared to laboratory mice and showed resilience against microbial changes (15). Moreover, we observed that WB-derived unique ASVs were more successful in colonizing in the Mix group piglets compared to the Sow inocula- derived ones. This colonization success of WB-derived ASV is perhaps due to the gut microbiota and host co-evolution over a long period leading to a mutualistic symbiosis where these microbes contribute to many of the host physiological processes, and the host provides a favorable niche for the microbes to colonize (59). A recent mice study of co-inoculation of native and non-native gut microbiota transplantation into germ-free mice revealed native microbiota receive adaptive advantage from the host and outcompete non-native microbiota (57). Another co-inoculation experiment reported that environmentally recruited co-adapted ‘native’ Burkholderia symbionts outcompete non-native bacteria in the gut of their bean bug host, even though they can colonize in the absence of the ‘native’ symbiont (60), which might explain why non-native Sow inoculum derived unique ASVs colonization into the Sow group piglets. These observations collectively suggest that wild boar gut-associated microbiota can effectively engraft and establish a stable microbial community in conventional piglets.

We also found that microbial richness in fecal and cecal samples of the Mix group was lower compared to the Control and WB groups, despite the Mix group having a higher initial microbial richness in the inoculum. This phenomenon likely occurred because of the introduction of two complex microbial communities into an already established gut environment that led to excessive competition during coalescence for niches and resources (61–63). As a result, many introduced microbes were unable to thrive and were subsequently eliminated. This competitive environment contributed to the Mix group’s cecal microbial community being distinct from the other groups. Despite the high competition, a subset of native microbes was able to colonize the Mix group successfully. Consequently, we observed that the fecal microbial community structure in the Mix group more closely resembled that of the WB and Sow groups, suggesting that inoculum-derived microbiota shaped the microbial composition.

The comparison of differential abundance of fecal taxa on day 6 post-transplantation revealed significant influences of the inoculum on gut microbiota composition despite the groups being housed under similar conditions. Inoculum-recipient piglets were enriched with bacterial taxa known for fiber degradation and SCFAs productions compared to the Control piglets. These findings indicate that microbial transplantation accelerated microbial metabolic maturation (64). In fact, a previous study reported that piglets exposed to soil were enriched with similar functional capabilities and became stable and mature gut microbiomes (65). Interestingly, on day 28 post-transplantation, we observed an enrichment of bacterial taxa across all groups capable of breaking down complex carbohydrates and producing SCFAs. Specifically, the WB group was enriched with taxa such as *Rikenellaceae RC9 gut group*, *Lactobacillus amylovorus*, *Veillonellaceae*, *Ruminococcaceae*, *Prevotellaceae UCG-001*, and *Gastranaerophilales* (66–70). In the Control group, there was an enrichment of *Prevotellaceae NK3B31*, *Oscillospirales UCG- 002*, and *Roseburia*. The Sow group showed an increase in *Megasphaera elsdenii* and *Lachnospiraceae_ND3007*. The Mix group had higher levels of *Lachnospiraceae NK4A136 group*, *Selenomonadaceae*, *Bacteroidales*, and *Oscillospirales UCG-010*. This is perhaps why we did not observe significant differences among treatment groups’ acetate, propionate, and butyrate levels. However, the WB group showed higher valeric acid levels than the Mix group.

Spearman correlation analysis between cecal valeric acid and bacterial taxa did not reveal any specific associations, but this difference may be attributed to the presence of *Lactobacillus* in the WB piglets, as many species within this genus are known to be associated with valeric acid production and confer beneficial outcomes for the host (71–73). Additionally, the WB piglets showed enrichment of *Gastranaerophilales*, a group from the Cyanobacteria phylum. Recent findings from our group also indicated that wild boars have a higher abundance of Cyanobacteria compared to their domestic counterparts (74). This could be due to their foraging in various environments, such as forests, grasslands, and wetlands, where they consume various plant materials, including roots, tubers, leaves, and seeds (75–77). This exposure potentially leads to the ingestion of soil, water, and plant material containing cyanobacteria. Intriguingly, cyanobacteria and their non-photosynthetic relatives are capable of participating in fermentation processes, breaking down complex polysaccharides, and producing SCFAs (77, 78), which could contribute to a mutually beneficial relationship within the host gut microbiome.

Beyond specific bacterial strain colonization, the success of FMT is also linked to the metabolic capacities of the engrafted microbes. Previous studies in human FMT have reported the importance of colonization of specific bacterial strains with metabolic capacity to produce beneficial metabolites for long-term success (38, 39). In our study, the cecal digesta in the WB group was enriched with histamine, acetyl-ornithine, tyramine, ornithine, citrulline, lactate, and glutamate. Histamine acts as a neurotransmitter for the “microbiota-gut-brain” axis and facilitates adverse conditions such as allergy reactions (79). Fecal scores, skin colour and extremities, body condition, and respiration rate throughout the experimental period were documented. However, we did not observe any such detrimental effect on the WB group.

Importantly, the mean weight gain also did not differ among the treatment groups. Intriguingly, studies demonstrated that gut commensal bacteria such as *Lactobacillus* can biosynthesize histamine and related products at normal physiological conditions (80, 81) and can play an anti-inflammatory role by controlling pro-inflammatory cytokine/chemokine production in the intestine (82). Interestingly, the WB inoculum was dominated by the *Lactobacillaceae* family, and three *Lactobacillus* taxa were found differentially abundant in the WB group cecal digesta compared to other groups. Correlative analysis between cecal microbiota and metabolites revealed that the histamine level was associated with *Lactobacillus* abundance. Interestingly, the precursor of histamine, histidine, was also found lower in the WB group and negatively correlated with *Lactobacillus* abundance. We also observed that the WB group had lower levels of IL-1β compared to the Control group and lower IL-6 levels compared to the Sow piglets. Studies have shown that different *Lactobacillus* strains that possess the *hdc* gene cluster which can reduce colitis through histamine H2 receptor-mediated IL-1β and IL-6 reduction in distal small intestine and cecum of mice (83–85). We also observed that histamine negatively correlated with the *SMB-53* genus. A recent study has reported that fermentable carbohydrate supplementation reduced cecal *SMB-53* and increased histamine levels in irritable bowel syndrome (IBS) mouse model (86). However, the role of luminal histamines under normal physiological conditions in the WB group is yet to be determined.

Like histamine, *Lactobacillus* abundance was positively associated with ornithine, acetyl- ornithine, citrulline, and glutamic acid. These metabolites are the product of the arginine deiminase (ADI) pathway, a metabolic trait observed most abundantly among *Lactobacillales* (87, 88). Citrulline supplementation has been associated with improved intestinal integrity in pig and mouse colitis and gut injury models (89–92). Moreover, studies have shown that *Lactobacillus*-derived L-ornithine increases L-kynurenine (a metabolite of tryptophan and a ligand for the AhR in intestinal epithelial cells). This L-kynurenine subsequently promotes the accumulation of RORγt(+)IL-22(+)ILC3 cells in gut tissues and maintains healthy gut mucosa (93). Similarly, glutamate supplementation in post-weaning piglets was associated with improved growth performance and gut health (94). Another study reported dietary glutamate reduced ileal IL-1β and IL-6, and improved growth, nutrient digestibility and increased tight junction protein expression (95).

Interestingly, *Lactobacillus* can utilize acetyl-ornithine from the ADI pathway to produce polyamines such as spermine, tyramine, and putrescine (81). This metabolic capability may explain the higher levels of tyramine and spermine observed in the WB group. Polyamines are important for the growth and maturation of the small intestine, which ameliorates intestinal atrophy by suppressing inflammatory cytokine including IL-6 in weaned piglets (96–98).

Spermine has been found to enhance antioxidant capacity and resistance to oxidative stress in piglets (99). Additionally, piglets on a diet supplemented with spermine and spermidine displayed increased growth compared to those on a standard diet (97). Cecal digesta of the Mix group were enriched with indole acetic acid with lower levels of IFN-γ, IL-2, and IL-6 compared to the Sow group. Previous studies have reported that indole derivatives can reduce IL-6, contributing to the maintenance of gut health and the prevention of inflammatory diseases in mouse and pig models (100, 101).

We also observed that the WB group had higher serotonin and dopamine levels in their cecal digesta. Like histamine, serotonin and dopamine act as neurotransmitters that can influence gastrointestinal motility, stress response, immune modulation, and overall health (102–105). A recent study in pigs showed that *Lactobacillus amylovorus* from the pig gut can produce serotonin through upregulating tryptophan hydroxylase 1 (*TPH1)* (106). Moreover, studies in mice and humans showed that *Lactobacillus*-derived serotonin not only regulates gut motility but also attenuates stress and depression through the microbiota-gut-brain axis (107, 108).

Interestingly, a study in rodents reported that dopamine can prepare the brain for heightened alertness, enabling it to shift attention toward potential threats (109). Therefore, the higher serotonin and dopamine levels in the WB inoculum recipient group may reflect a service provided by co-evolved microbes.

Despite these promising findings, several limitations in this study necessitate a cautious interpretation of the findings. To validate the observed changes in gut microbiota and metabolites, and to assess the impact of FMT on growth performance, large-scale experiments with more replicates under varied environmental conditions are necessary. Additionally, the study employed *16S rRNA* gene-based amplicon sequencing, which lacks the resolution to identify microbes at the strain level and does not provide functional profiling. To overcome these limitations, future studies incorporating a high-resolution metagenomic approach are warranted to accurately resolve taxonomic assignments and assess the functional capacity of microbial communities.

In conclusion, our study revealed that a successful colonization of wild boar-derived microbes to domestic pigs early in life via cultured MMC can alter gut microbial communities as well as cecal metabolite and cytokine profiles without causing adverse effects. Further research is needed to explore the relationship between these microbial-associated metabolites and immune modulation. Additionally, infection challenge studies are warranted to evaluate the disease resistance potential of these microbial communities.

## Acknowledgements

This study received funding from the Alberta Livestock and Meat Agency (res0030386) and a Discovery grant from the Natural Sciences and Engineering Research Council of Canada (RGPIN-2019–06336). B.P.W. was supported through the Canada Research Chair Program, while the Alberta Graduate Excellence Scholarship and the Frank Aherne Graduate Scholarship in Swine Research supported R.R. We extend our gratitude to the staff at the Animal Care Unit of the Western College of Veterinary Medicine at the University of Saskatchewan for their invaluable guidance and support. We also acknowledge the funding from the University of Saskatchewan and the United States Animal and Plant Health Inspection Service National Feral Swine Damage Management Program for facilitating the capture of wild pigs.

## Authors’ contributions

B.P.W. conceived the study and secured the funding. R.R. and J.M.F. designed and conducted the experiment with assistance from R.K.B., R.N., and J.H. R.R. and J.M.F. also performed the laboratory work. Bioinformatic and statistical analyses were carried out by R.R., J.M.F., Y.F., T.B., and T.J. R.R. wrote and revised the main manuscript, incorporating input from all authors. All authors read and approved the final manuscript.

## Data availability

The raw 16S rRNA gene amplicon sequence read data is available through the National Center for Biotechnology Information (NCBI) Sequence Read Archive (BioProject Accession number PRJNA1179462). Cecal metabolomics data can be obtained upon request from the corresponding

---

**Supplementary Figure 1.**
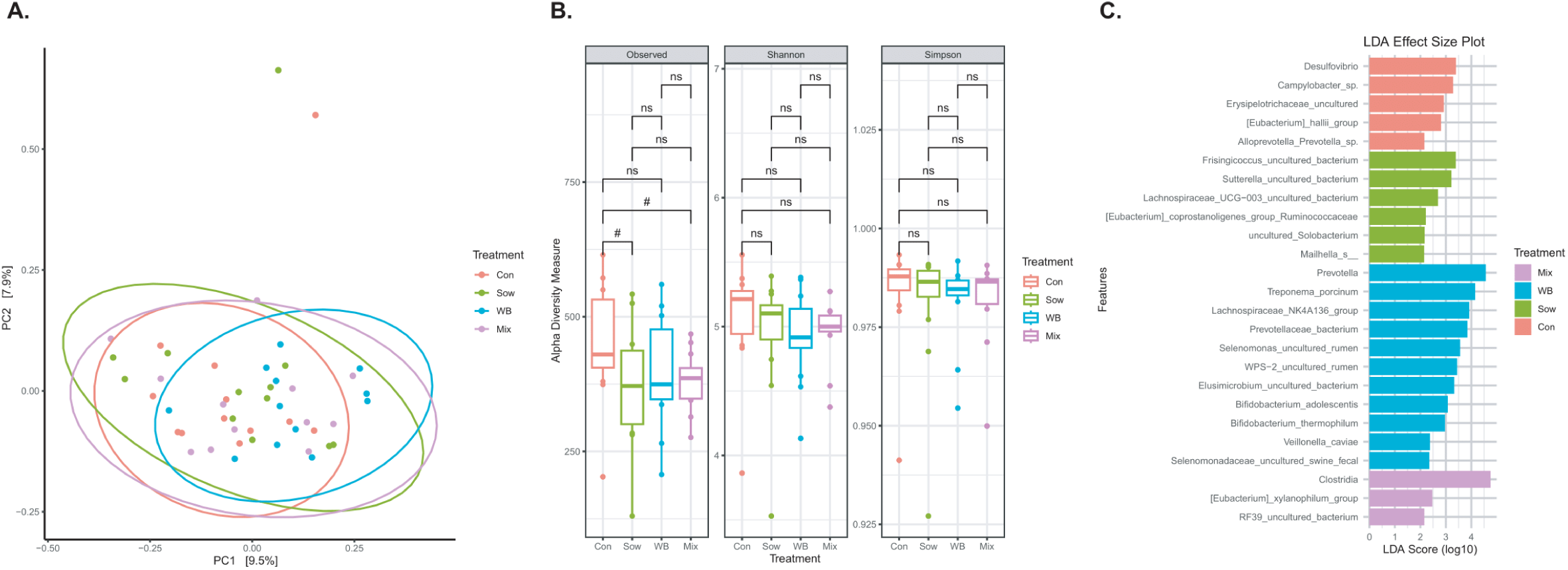
Comparison of post-transplantation day 6 (PND 27) fecal microbiota composition and α-diversity among treatment groups (n = 12/group). (A) At day 6 (PND 27) post-transplantation, there was a trend of a shift in microbial community structure as measured by β-diversity based on Bray–Curtis dissimilarity (Adonis, R² = 0.07, *P* = 0.07; Betadispersion *P* = 0.90). (B) No differences were observed in α--diversity indices among treatment groups at day 6 (PND 27) post-transplantation (Observed *P* = 0.18; Shannon *P* = 0.36; Simpson *P* = 0.38) (C) LEfSe analysis identified differentially abundant taxonomical features among treatment groups (*P* < 0.05). \**P* < 0.05, \*\**P* < 0.01, \*\*\**P* < 0.001, *^#^P* > 0.05 and < 0.1

**Supplementary Figure 2:**
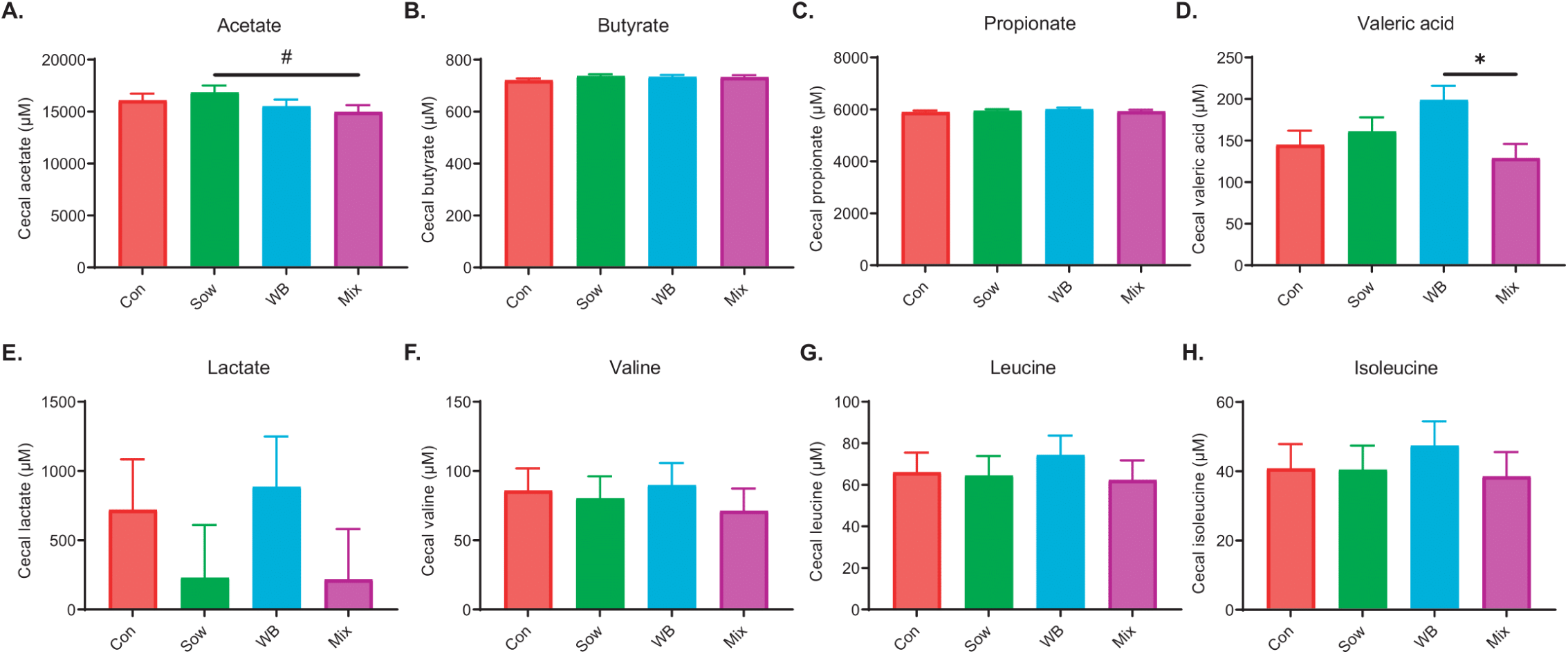
Comparison of cecal SCFAs, lactate, and branch-chain amino acid concentrations among treatment groups. (A) Acetate; (B) Butyrate; (C) Propionate; (D) Valeric acid; (E) Lactate (F) Valine; (G) Leucine; and (H) Isoleucine (n = 12/group). All data is shown as a mean with SEM. *P*-values were obtained from adjusted pairwise comparisons after a linear mixed effect model (where litter and pen effect accounted for a random effect) and significance was considered at *P <* 0.05. \**P* < 0.05, \*\**P* < 0.01, \*\*\**P* < 0.001, *^#^P* > 0.05 and < 0.1

**Supplementary Figure 3:**
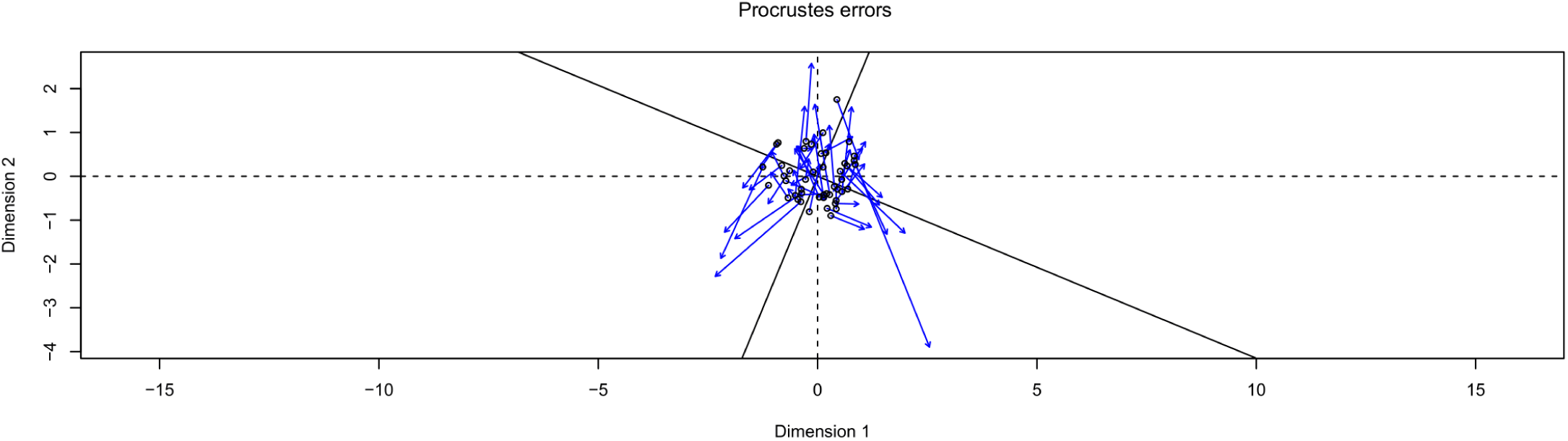
Procrustes analysis on the principal-coordinate analysis axes of the cecal microbiome and metabolome. An association between cecal metabolites and microbiota compositions was observed (Procrustes sum of squares, 0.75; correlation, 0.49; *P* < 0.001). Greater distances between the black hollow circles representing metabolite eigenvalues and the blue arrows representing microbiota eigenvalues indicated a higher level of discordance between the datasets for each sample.

**Supplementary Table 1:** List of 70 metabolites in TMIC prime assay.

| *Metabolite class* | *Metabolite* |
| --- | --- |
| *Amine Oxide* | Trimethylamine N-oxide |
| *Amino Acids* | Alanine, Arginine, Asparagine, Aspartate, Citrulline, Glutamine, Glutamate, Glycine, Histidine, Isoleucine, Leucine, Lysine, Methionine, Ornithine, Phenylalanine, Proline, Serine, Threonine, Tryptophan, Tyrosine, Valine, Betaine |
| *Amino Acid Derivatives* | Creatine, Phosphocreatine, Methylhistidine |
| *Biogenic Amines* | Acetyl-ornithine, ADMA, Asymmetric dimethylarginine, Total dimethylarginine, Alpha-Aminoadipic acid, Carnosine, Creatinine, Dihydroxyphenylalanine, Dopamine, Histamine, Kynurenine, Methioninesulfoxide, Hydroxyproline, Hydroxyproline, Nitrotyrosine, Phenylethylamine, Putrescine, Sarcosine, Serotonin, Spermidine, Spermine, Diacetylspermine, Taurine, Tyramine |
| *Monosaccharides* | Glucose |
| *Organic Acids* | Lactate, Beta-hydroxybutyric acid, Alpha-ketoglutarate, Citric acid, Butyrate, Propionic acid, Valeric acid, Para-hydroxyhippuric acid, Succinic acid, Fumaric acid, Pyruvic acid, Isobutyric acid, Hippuric acid, Methylmalonic acid, Acetic acid, Indole acetic acid, Uric acid |
| *Vitamins & Cofactors* | Choline |

ADMA: Asymmetric dimethylarginine

## References

1. Stappenbeck TS, Virgin HW. 2016. Accounting for reciprocal host-microbiome interactions in experimental science. Nature 534:191–9.

2. Sommer F, Backhed F. 2013. The gut microbiota--masters of host development and physiology. Nat Rev Microbiol 11:227–38.

3. Henry LP, Bruijning M, Forsberg SKG, Ayroles JF. 2021. The microbiome extends host evolutionary potential. Nat Commun 12:5141.

4. Lee WJ, Hase K. 2014. Gut microbiota-generated metabolites in animal health and disease. Nat Chem Biol 10:416–24.

5. Hou K, Wu ZX, Chen XY, Wang JQ, Zhang D, Xiao C, Zhu D, Koya JB, Wei L, Li J, Chen ZS. 2022. Microbiota in health and diseases. Signal Transduct Target Ther 7:135.

6. Guevarra RB, Lee JH, Lee SH, Seok MJ, Kim DW, Kang BN, Johnson TJ, Isaacson RE, Kim HB. 2019. Piglet gut microbial shifts early in life: causes and effects. J Anim Sci Biotechnol 10:1.

7. Gensollen T, Iyer SS, Kasper DL, Blumberg RS. 2016. How colonization by microbiota in early life shapes the immune system. Science 352:539–44.

8. Rahman R, Fouhse JM, Prisnee TL, Ju T, Diether NE, Willing BP. 2023. Comparing the impact of mixed-culture microbial communities and fecal transplant on the intestinal microbiota and metabolome of weaned piglets. FEMS Microbiol Ecol 99.

9. Te Pas MFW, Jansman AJM, Kruijt L, van der Meer Y, Vervoort JJM, Schokker D. 2020. Sanitary Conditions Affect the Colonic Microbiome and the Colonic and Systemic Metabolome of Female Pigs. Front Vet Sci 7:585730.

10. Law K, Lozinski B, Torres I, Davison S, Hilbrands A, Nelson E, Parra-Suescun J, Johnston L, Gomez A. 2021. Disinfection of Maternal Environments Is Associated with Piglet Microbiome Composition from Birth to Weaning. mSphere 6:e0066321.

11. Kuthyar S, Diaz J, Avalos-Villatoro F, Maltecca C, Tiezzi F, Dunn RR, Reese AT. 2023. Domestication shapes the pig gut microbiome and immune traits from the scale of lineage to population. J Evol Biol 36:1695–1711.

12. Zhang S, Zhang H, Zhang C, Wang G, Shi C, Li Z, Gao F, Cui Y, Li M, Yang G. 2024. Composition and evolutionary characterization of the gut microbiota in pigs. Int Microbiol 27:993–1008.

13. Wei L, Zhou W, Zhu Z. 2022. Comparison of Changes in Gut Microbiota in Wild Boars and Domestic Pigs Using 16S rRNA Gene and Metagenomics Sequencing Technologies. Animals (Basel) 12.

14. Petrelli S, Buglione M, Rivieccio E, Ricca E, Baccigalupi L, Scala G, Fulgione D. 2023. Reprogramming of the gut microbiota following feralization in Sus scrofa. Anim Microbiome 5:14.

15. Rosshart SP, Herz J, Vassallo BG, Hunter A, Wall MK, Badger JH, McCulloch JA, Anastasakis DG, Sarshad AA, Leonardi I, Collins N, Blatter JA, Han SJ, Tamoutounour S, Potapova S, Foster St Claire MB, Yuan W, Sen SK, Dreier MS, Hild B, Hafner M, Wang D, Iliev ID, Belkaid Y, Trinchieri G, Rehermann B. 2019. Laboratory mice born to wild mice have natural microbiota and model human immune responses. Science 365.

16. Rosshart SP, Vassallo BG, Angeletti D, Hutchinson DS, Morgan AP, Takeda K, Hickman HD, McCulloch JA, Badger JH, Ajami NJ, Trinchieri G, Pardo-Manuel de Villena F, Yewdell JW, Rehermann B. 2017. Wild Mouse Gut Microbiota Promotes Host Fitness and Improves Disease Resistance. Cell 171:1015–1028 e13.

17. Hu J, Chen J, Ma L, Hou Q, Zhang Y, Kong X, Huang X, Tang Z, Wei H, Wang X, Yan X. 2024. Characterizing core microbiota and regulatory functions of the pig gut microbiome. ISME J 18.

18. Gresse R, Chaucheyras-Durand F, Fleury MA, Van de Wiele T, Forano E, Blanquet-Diot S. 2017. Gut Microbiota Dysbiosis in Postweaning Piglets: Understanding the Keys to Health. Trends Microbiol 25:851–873.

19. Chen L, Xu Y, Chen X, Fang C, Zhao L, Chen F. 2017. The Maturing Development of Gut Microbiota in Commercial Piglets during the Weaning Transition. Front Microbiol 8:1688.

20. Gebhardt JT, Tokach MD, Dritz SS, DeRouchey JM, Woodworth JC, Goodband RD, Henry SC. 2020. Postweaning mortality in commercial swine production. I: review of non-infectious contributing factors. Transl Anim Sci 4:txaa068.

21. Looft T, Allen HK, Cantarel BL, Levine UY, Bayles DO, Alt DP, Henrissat B, Stanton TB. 2014. Bacteria, phages and pigs: the effects of in-feed antibiotics on the microbiome at different gut locations. ISME J 8:1566–76.

22. Kim K, Jinno C, Ji P, Liu Y. 2022. Trace amounts of antibiotic altered metabolomic and microbial profiles of weaned pigs infected with a pathogenic E. coli. J Anim Sci Biotechnol 13:59.

23. Roca I, Akova M, Baquero F, Carlet J, Cavaleri M, Coenen S, Cohen J, Findlay D, Gyssens I, Heuer OE, Kahlmeter G, Kruse H, Laxminarayan R, Liebana E, Lopez-Cerero L, MacGowan A, Martins M, Rodriguez-Bano J, Rolain JM, Segovia C, Sigauque B, Tacconelli E, Wellington E, Vila J. 2015. The global threat of antimicrobial resistance: science for intervention. New Microbes New Infect 6:22–9.

24. Hoffmann DE, Palumbo FB, Ravel J, Rowthorn V, von Rosenvinge E. 2017. A proposed definition of microbiota transplantation for regulatory purposes. Gut Microbes 8:208–213.

25. Quraishi MN, Widlak M, Bhala N, Moore D, Price M, Sharma N, Iqbal TH. 2017. Systematic review with meta-analysis: the efficacy of faecal microbiota transplantation for the treatment of recurrent and refractory Clostridium difficile infection. Aliment Pharmacol Ther 46:479–493.

26. Moayyedi P, Surette MG, Kim PT, Libertucci J, Wolfe M, Onischi C, Armstrong D, Marshall JK, Kassam Z, Reinisch W, Lee CH. 2015. Fecal Microbiota Transplantation Induces Remission in Patients With Active Ulcerative Colitis in a Randomized Controlled Trial. Gastroenterology 149:102–109 e6.

27. Paramsothy S, Kamm MA, Kaakoush NO, Walsh AJ, van den Bogaerde J, Samuel D, Leong RWL, Connor S, Ng W, Paramsothy R, Xuan W, Lin E, Mitchell HM, Borody TJ. 2017. Multidonor intensive faecal microbiota transplantation for active ulcerative colitis: a randomised placebo-controlled trial. Lancet 389:1218–1228.

28. Zhang J, Rodriguez F, Navas MJ, Costa-Hurtado M, Almagro V, Bosch-Camos L, Lopez E, Cuadrado R, Accensi F, Pina-Pedrero S, Martinez J, Correa-Fiz F. 2020. Fecal microbiota transplantation from warthog to pig confirms the influence of the gut microbiota on African swine fever susceptibility. Sci Rep 10:17605.

29. Xiang Q, Wu X, Pan Y, Wang L, Cui C, Guo Y, Zhu L, Peng J, Wei H. 2020. Early-Life Intervention Using Fecal Microbiota Combined with Probiotics Promotes Gut Microbiota Maturation, Regulates Immune System Development, and Alleviates Weaning Stress in Piglets. Int J Mol Sci 21.

30. Brunse A, Martin L, Rasmussen TS, Christensen L, Skovsted Cilieborg M, Wiese M, Khakimov B, Pieper R, Nielsen DS, Sangild PT, Thymann T. 2019. Effect of fecal microbiota transplantation route of administration on gut colonization and host response in preterm pigs. ISME J 13:720–733.

31. McCormack UM, Curiao T, Wilkinson T, Metzler-Zebeli BU, Reyer H, Ryan T, Calderon-Diaz JA, Crispie F, Cotter PD, Creevey CJ, Gardiner GE, Lawlor PG. 2018. Fecal Microbiota Transplantation in Gestating Sows and Neonatal Offspring Alters Lifetime Intestinal Microbiota and Growth in Offspring. mSystems 3.

32. Qi R, Zhang Z, Wang J, Qiu X, Wang Q, Yang F, Huang J, Liu Z. 2021. Introduction of Colonic and Fecal Microbiota From an Adult Pig Differently Affects the Growth, Gut Health, Intestinal Microbiota and Blood Metabolome of Newborn Piglets. Front Microbiol 12:623673.

33. Zuo T, Wong SH, Cheung CP, Lam K, Lui R, Cheung K, Zhang F, Tang W, Ching JYL, Wu JCY, Chan PKS, Sung JJY, Yu J, Chan FKL, Ng SC. 2018. Gut fungal dysbiosis correlates with reduced efficacy of fecal microbiota transplantation in Clostridium difficile infection. Nat Commun 9:3663.

34. Park H, Laffin MR, Jovel J, Millan B, Hyun JE, Hotte N, Kao D, Madsen KL. 2019. The success of fecal microbial transplantation in Clostridium difficile infection correlates with bacteriophage relative abundance in the donor: a retrospective cohort study. Gut Microbes 10:676–687.

35. Paramsothy S, Nielsen S, Kamm MA, Deshpande NP, Faith JJ, Clemente JC, Paramsothy R, Walsh AJ, van den Bogaerde J, Samuel D, Leong RWL, Connor S, Ng W, Lin E, Borody TJ, Wilkins MR, Colombel JF, Mitchell HM, Kaakoush NO. 2019. Specific Bacteria and Metabolites Associated With Response to Fecal Microbiota Transplantation in Patients With Ulcerative Colitis. Gastroenterology 156:1440–1454 e2.

36. Kump P, Wurm P, Grochenig HP, Wenzl H, Petritsch W, Halwachs B, Wagner M, Stadlbauer V, Eherer A, Hoffmann KM, Deutschmann A, Reicht G, Reiter L, Slawitsch P, Gorkiewicz G, Hogenauer C. 2018. The taxonomic composition of the donor intestinal microbiota is a major factor influencing the efficacy of faecal microbiota transplantation in therapy refractory ulcerative colitis. Aliment Pharmacol Ther 47:67–77.

37. Jiang ZD, Ajami NJ, Petrosino JF, Jun G, Hanis CL, Shah M, Hochman L, Ankoma-Sey V, DuPont AW, Wong MC, Alexander A, Ke S, DuPont HL. 2017. Randomised clinical trial: faecal microbiota transplantation for recurrent Clostridum difficile infection - fresh, or frozen, or lyophilised microbiota from a small pool of healthy donors delivered by colonoscopy. Aliment Pharmacol Ther 45:899–908.

38. Watson AR, Fussel J, Veseli I, DeLongchamp JZ, Silva M, Trigodet F, Lolans K, Shaiber A, Fogarty E, Runde JM, Quince C, Yu MK, Soylev A, Morrison HG, Lee STM, Kao D, Rubin DT, Jabri B, Louie T, Eren AM. 2023. Metabolic independence drives gut microbial colonization and resilience in health and disease. Genome Biol 24:78.

39. Podlesny D, Durdevic M, Paramsothy S, Kaakoush NO, Hogenauer C, Gorkiewicz G, Walter J, Fricke WF. 2022. Identification of clinical and ecological determinants of strain engraftment after fecal microbiota transplantation using metagenomics. Cell Rep Med 3:100711.

40. Patil Y, Gooneratne R, Ju XH. 2020. Interactions between host and gut microbiota in domestic pigs: a review. Gut Microbes 11:310–334.

41. Megahed A, Zeineldin M, Evans K, Maradiaga N, Blair B, Aldridge B, Lowe J. 2019. Impacts of environmental complexity on respiratory and gut microbiome community structure and diversity in growing pigs. Sci Rep 9:13773.

42. Ericsson AC, Gagliardi J, Bouhan D, Spollen WG, Givan SA, Franklin CL. 2018. The influence of caging, bedding, and diet on the composition of the microbiota in different regions of the mouse gut. Sci Rep 8:4065.

43. Barbosa JA, Rodrigues LA, Columbus DA, Aguirre JCP, Harding JCS, Cantarelli VS, Costa MO. 2021. Experimental infectious challenge in pigs leads to elevated fecal calprotectin levels following colitis, but not enteritis. Porcine Health Manag 7:48.

44. Willing BP, Vacharaksa A, Croxen M, Thanachayanont T, Finlay BB. 2011. Altering host resistance to infections through microbial transplantation. PLoS One 6:e26988.

45. Bolyen E, Rideout JR, Dillon MR, Bokulich NA, Abnet CC, Al-Ghalith GA, Alexander H, Alm EJ, Arumugam M, Asnicar F, Bai Y, Bisanz JE, Bittinger K, Brejnrod A, Brislawn CJ, Brown CT, Callahan BJ, Caraballo-Rodriguez AM, Chase J, Cope EK, Da Silva R, Diener C, Dorrestein PC, Douglas GM, Durall DM, Duvallet C, Edwardson CF, Ernst M, Estaki M, Fouquier J, Gauglitz JM, Gibbons SM, Gibson DL, Gonzalez A, Gorlick K, Guo J, Hillmann B, Holmes S, Holste H, Huttenhower C, Huttley GA, Janssen S, Jarmusch AK, Jiang L, Kaehler BD, Kang KB, Keefe CR, Keim P, Kelley ST, Knights D, et al. 2019. Reproducible, interactive, scalable and extensible microbiome data science using QIIME 2. Nat Biotechnol 37:852–857.

46. Callahan BJ, McMurdie PJ, Rosen MJ, Han AW, Johnson AJ, Holmes SP. 2016. DADA2: High-resolution sample inference from Illumina amplicon data. Nat Methods 13:581–3.

47. Katoh K, Misawa K, Kuma K, Miyata T. 2002. MAFFT: a novel method for rapid multiple sequence alignment based on fast Fourier transform. Nucleic Acids Res 30:3059–66.

48. Bokulich NA, Kaehler BD, Rideout JR, Dillon M, Bolyen E, Knight R, Huttley GA, Gregory Caporaso J. 2018. Optimizing taxonomic classification of marker-gene amplicon sequences with QIIME 2’s q2-feature-classifier plugin. Microbiome 6:90.

49. Pedregosa F, Varoquaux G, Gramfort A, Michel V, Thirion B, Grisel O, Blondel M, Prettenhofer P, Weiss R, Dubourg V. 2011. Scikit-learn: Machine learning in Python. the Journal of machine Learning research 12:2825–2830.

50. Quast C, Pruesse E, Yilmaz P, Gerken J, Schweer T, Yarza P, Peplies J, Glockner FO. 2013. The SILVA ribosomal RNA gene database project: improved data processing and web-based tools. Nucleic Acids Res 41:D590–6.

51. McMurdie PJ, Holmes S. 2013. phyloseq: an R package for reproducible interactive analysis and graphics of microbiome census data. PLoS One 8:e61217.

52. Moeller AH, Caro-Quintero A, Mjungu D, Georgiev AV, Lonsdorf EV, Muller MN, Pusey AE, Peeters M, Hahn BH, Ochman H. 2016. Cospeciation of gut microbiota with hominids. Science 353:380–2.

53. Reese AT, Chadaideh KS, Diggins CE, Schell LD, Beckel M, Callahan P, Ryan R, Emery Thompson M, Carmody RN. 2021. Effects of domestication on the gut microbiota parallel those of human industrialization. Elife 10.

54. Ang L, Vinderola G, Endo A, Kantanen J, Jingfeng C, Binetti A, Burns P, Qingmiao S, Suying D, Zujiang Y, Rios-Covian D, Mantziari A, Beasley S, Gomez-Gallego C, Gueimonde M, Salminen S. 2022. Gut Microbiome Characteristics in feral and domesticated horses from different geographic locations. Commun Biol 5:172.

55. Sidiropoulos DN, Al-Ghalith GA, Shields-Cutler RR, Ward TL, Johnson AJ, Vangay P, Knights D, Kashyap PC, Xian Y, Ramer-Tait AE, Clayton JB. 2020. Wild primate microbiomes prevent weight gain in germ-free mice. Anim Microbiome 2:16.

56. Prabhu VR, Wasimuddin, Kamalakkannan R, Arjun MS, Nagarajan M. 2020. Consequences of Domestication on Gut Microbiome: A Comparative Study Between Wild Gaur and Domestic Mithun. Front Microbiol 11:133.

57. Sprockett DD, Price JD, Juritsch AF, Schmaltz RJ, Real MVF, Goldman SL, Sheehan M, Ramer-Tait AE, Moeller AH. 2023. Home-site advantage for host species-specific gut microbiota. Sci Adv 9:eadf5499.

58. Ley RE, Peterson DA, Gordon JI. 2006. Ecological and evolutionary forces shaping microbial diversity in the human intestine. Cell 124:837–48.

59. Larzul C, Estelle J, Borey M, Blanc F, Lemonnier G, Billon Y, Thiam MG, Quinquis B, Galleron N, Jardet D, Lecardonnel J, Plaza Onate F, Rogel-Gaillard C. 2024. Driving gut microbiota enterotypes through host genetics. Microbiome 12:116.

60. Itoh H, Jang S, Takeshita K, Ohbayashi T, Ohnishi N, Meng XY, Mitani Y, Kikuchi Y. 2019. Host-symbiont specificity determined by microbe-microbe competition in an insect gut. Proc Natl Acad Sci U S A 116:22673–22682.

61. Lechon-Alonso P, Clegg T, Cook J, Smith TP, Pawar S. 2021. The role of competition versus cooperation in microbial community coalescence. PLoS Comput Biol 17:e1009584.

62. Costello EK, Stagaman K, Dethlefsen L, Bohannan BJ, Relman DA. 2012. The application of ecological theory toward an understanding of the human microbiome. Science 336:1255–62.

63. Keshteli AH, Millan B, Madsen KL. 2017. Pretreatment with antibiotics may enhance the efficacy of fecal microbiota transplantation in ulcerative colitis: a meta-analysis. Mucosal Immunol 10:565–566.

64. Wang X, Tsai T, Deng F, Wei X, Chai J, Knapp J, Apple J, Maxwell CV, Lee JA, Li Y, Zhao J. 2019. Longitudinal investigation of the swine gut microbiome from birth to market reveals stage and growth performance associated bacteria. Microbiome 7:109.

65. Vo N, Tsai TC, Maxwell C, Carbonero F. 2017. Early exposure to agricultural soil accelerates the maturation of the early-life pig gut microbiota. Anaerobe 45:31–39.

66. Fusco W, Lorenzo MB, Cintoni M, Porcari S, Rinninella E, Kaitsas F, Lener E, Mele MC, Gasbarrini A, Collado MC, Cammarota G, Ianiro G. 2023. Short-Chain Fatty-Acid-Producing Bacteria: Key Components of the Human Gut Microbiota. Nutrients 15.

67. Deleu S, Machiels K, Raes J, Verbeke K, Vermeire S. 2021. Short chain fatty acids and its producing organisms: An overlooked therapy for IBD? EBioMedicine 66:103293.

68. Qiu M, Hu J, Peng H, Li B, Xu J, Song X, Yu C, Zhang Z, Du X, Bu G, Huang A, Han X, Zeng X, Yang C, Kong F. 2022. Research Note: The gut microbiota varies with dietary fiber levels in broilers. Poult Sci 101:101922.

69. Flint HJ, Duncan SH, Scott KP, Louis P. 2015. Links between diet, gut microbiota composition and gut metabolism. Proc Nutr Soc 74:13–22.

70. Morrison DJ, Preston T. 2016. Formation of short chain fatty acids by the gut microbiota and their impact on human metabolism. Gut Microbes 7:189–200.

71. Lau HC, Zhang X, Ji F, Lin Y, Liang W, Li Q, Chen D, Fong W, Kang X, Liu W, Chu ES, Ng QW, Yu J. 2024. Lactobacillus acidophilus suppresses non-alcoholic fatty liver disease-associated hepatocellular carcinoma through producing valeric acid. EBioMedicine 100:104952.

72. Lin X, Xiao HM, Liu HM, Lv WQ, Greenbaum J, Gong R, Zhang Q, Chen YC, Peng C, Xu XJ, Pan DY, Chen Z, Li ZF, Zhou R, Wang XF, Lu JM, Ao ZX, Song YQ, Zhang YH, Su KJ, Meng XH, Ge CL, Lv FY, Luo Z, Shi XM, Zhao Q, Guo BY, Yi NJ, Shen H, Papasian CJ, Shen J, Deng HW. 2023. Gut microbiota impacts bone via Bacteroides vulgatus-valeric acid-related pathways. Nat Commun 14:6853.

73. Hemalatha R, Ouwehand AC, Saarinen MT, Prasad UV, Swetha K, Bhaskar V. 2017. Effect of probiotic supplementation on total lactobacilli, bifidobacteria and short chain fatty acids in 2-5-year-old children. Microb Ecol Health Dis 28:1298340.

74. Rahman R, Fouhse JM, Ju T, Fan Y, C SM, Pieper R, Brook RK, Willing BP. 2024. A comparison of wild boar and domestic pig microbiota does not reveal a loss of microbial species but an increase in alpha diversity and opportunistic genera in domestic pigs. Microbiol Spectr doi:10.1128/spectrum.00843-24:e0084324.

75. McGorum BC, Pirie RS, Glendinning L, McLachlan G, Metcalf JS, Banack SA, Cox PA, Codd GA. 2015. Grazing livestock are exposed to terrestrial cyanobacteria. Vet Res 46:16.

76. Monchamp ME, Spaak P, Pomati F. 2018. Long Term Diversity and Distribution of Non-photosynthetic Cyanobacteria in Peri-Alpine Lakes. Front Microbiol 9:3344.

77. Di Rienzi SC, Sharon I, Wrighton KC, Koren O, Hug LA, Thomas BC, Goodrich JK, Bell JT, Spector TD, Banfield JF, Ley RE. 2013. The human gut and groundwater harbor non-photosynthetic bacteria belonging to a new candidate phylum sibling to Cyanobacteria. Elife 2:e01102.

78. Utami YD, Kuwahara H, Murakami T, Morikawa T, Sugaya K, Kihara K, Yuki M, Lo N, Deevong P, Hasin S, Boonriam W, Inoue T, Yamada A, Ohkuma M, Hongoh Y. 2018. Phylogenetic Diversity and Single-Cell Genome Analysis of “Melainabacteria”, a Non-Photosynthetic Cyanobacterial Group, in the Termite Gut. Microbes Environ 33:50–57.

79. Thurmond RL, Gelfand EW, Dunford PJ. 2008. The role of histamine H1 and H4 receptors in allergic inflammation: the search for new antihistamines. Nat Rev Drug Discov 7:41–53.

80. Beaver MH, Wostmann BS. 1962. Histamine and 5-hydroxytryptamine in the intestinal tract of germ-free animals, animals harbouring one microbial species and conventional animals. Br J Pharmacol Chemother 19:385–93.

81. Pugin B, Barcik W, Westermann P, Heider A, Wawrzyniak M, Hellings P, Akdis CA, O’Mahony L. 2017. A wide diversity of bacteria from the human gut produces and degrades biogenic amines. Microb Ecol Health Dis 28:1353881.

82. Levy M, Thaiss CA, Zeevi D, Dohnalova L, Zilberman-Schapira G, Mahdi JA, David E, Savidor A, Korem T, Herzig Y, Pevsner-Fischer M, Shapiro H, Christ A, Harmelin A, Halpern Z, Latz E, Flavell RA, Amit I, Segal E, Elinav E. 2015. Microbiota-Modulated Metabolites Shape the Intestinal Microenvironment by Regulating NLRP6 Inflammasome Signaling. Cell 163:1428–43.

83. Gao C, Major A, Rendon D, Lugo M, Jackson V, Shi Z, Mori-Akiyama Y, Versalovic J. 2015. Histamine H2 Receptor-Mediated Suppression of Intestinal Inflammation by Probiotic Lactobacillus reuteri. mBio 6:e01358–15.

84. Ganesh BP, Hall A, Ayyaswamy S, Nelson JW, Fultz R, Major A, Haag A, Esparza M, Lugo M, Venable S, Whary M, Fox JG, Versalovic J. 2018. Diacylglycerol kinase synthesized by commensal Lactobacillus reuteri diminishes protein kinase C phosphorylation and histamine-mediated signaling in the mammalian intestinal epithelium. Mucosal Immunol 11:380–393.

85. Kim J, Balasubramanian I, Bandyopadhyay S, Nadler I, Singh R, Harlan D, Bumber A, He Y, Kerkhof LJ, Gao N, Su X, Ferraris RP. 2021. Lactobacillus rhamnosus GG modifies the metabolome of pathobionts in gnotobiotic mice. BMC Microbiol 21:165.

86. De Palma G, Shimbori C, Reed DE, Yu Y, Rabbia V, Lu J, Jimenez-Vargas N, Sessenwein J, Lopez-Lopez C, Pigrau M, Jaramillo-Polanco J, Zhang Y, Baerg L, Manzar A, Pujo J, Bai X, Pinto-Sanchez MI, Caminero A, Madsen K, Surette MG, Beyak M, Lomax AE, Verdu EF, Collins SM, Vanner SJ, Bercik P. 2022. Histamine production by the gut microbiota induces visceral hyperalgesia through histamine 4 receptor signaling in mice. Sci Transl Med 14:eabj1895.

87. Noens EE, Lolkema JS. 2017. Convergent evolution of the arginine deiminase pathway: the ArcD and ArcE arginine/ornithine exchangers. Microbiologyopen 6.

88. Zuniga M, Perez G, Gonzalez-Candelas F. 2002. Evolution of arginine deiminase (ADI) pathway genes. Mol Phylogenet Evol 25:429–44.

89. Kvidera SK, Mayorga EJ, McCarthy CS, Horst EA, Abeyta MA, Baumgard LH. 2024. Effects of supplemental citrulline on thermal and intestinal morphology parameters during heat stress and feed restriction in growing pigs. J Anim Sci 102.

90. Chapman JC, Liu Y, Zhu L, Rhoads JM. 2012. Arginine and citrulline protect intestinal cell monolayer tight junctions from hypoxia-induced injury in piglets. Pediatr Res 72:576–82.

91. van Wijck K, Wijnands KA, Meesters DM, Boonen B, van Loon LJ, Buurman WA, Dejong CH, Lenaerts K, Poeze M. 2014. L-citrulline improves splanchnic perfusion and reduces gut injury during exercise. Med Sci Sports Exerc 46:2039–46.

92. Antunes MM, Leocadio PC, Teixeira LG, Leonel AJ, Cara DC, Menezes GB, Generoso Sde V, Cardoso VN, Alvarez-Leite JI, Correia MI. 2016. Pretreatment With L-Citrulline Positively Affects the Mucosal Architecture and Permeability of the Small Intestine in a Murine Mucositis Model. JPEN J Parenter Enteral Nutr 40:279–86.

93. Qi H, Li Y, Yun H, Zhang T, Huang Y, Zhou J, Yan H, Wei J, Liu Y, Zhang Z, Gao Y, Che Y, Su X, Zhu D, Zhang Y, Zhong J, Yang R. 2019. Lactobacillus maintains healthy gut mucosa by producing L-Ornithine. Commun Biol 2:171.

94. Luise D, Correa F, Chalvon-Demersay T, Galosi L, Rossi G, Lambert W, Bosi P, Trevisi P. 2022. Supplementation of mixed doses of glutamate and glutamine can improve the growth and gut health of piglets during the first 2 weeks post-weaning. Sci Rep 12:14533.

95. Kyoung H, Lee JJ, Cho JH, Choe J, Kang J, Lee H, Liu Y, Kim Y, Kim HB, Song M. 2021. Dietary Glutamic Acid Modulates Immune Responses and Gut Health of Weaned Pigs. Animals (Basel) 11.

96. Sabater-Molina M, Larque E, Torrella F, Plaza J, Lozano T, Munoz A, Zamora S. 2009. Effects of dietary polyamines at physiologic doses in early-weaned piglets. Nutrition 25:940–6.

97. van Wettere WH, Willson NL, Pain SJ, Forder RE. 2016. Effect of oral polyamine supplementation pre-weaning on piglet growth and intestinal characteristics. Animal 10:1655–9.

98. Liu B, Jiang X, Cai L, Zhao X, Dai Z, Wu G, Li X. 2019. Putrescine mitigates intestinal atrophy through suppressing inflammatory response in weanling piglets. J Anim Sci Biotechnol 10:69.

99. Fang T, Zheng J, Cao W, Jia G, Zhao H, Chen X, Cai J, Wang J, Liu G. 2018. Effects of spermine on the antioxidant status and gene expression of antioxidant-related signaling molecules in the liver and longissimus dorsi of piglets. Animal 12:1208–1216.

100. Shen J, Yang L, You K, Chen T, Su Z, Cui Z, Wang M, Zhang W, Liu B, Zhou K, Lu H. 2022. Indole-3-Acetic Acid Alters Intestinal Microbiota and Alleviates Ankylosing Spondylitis in Mice. Front Immunol 13:762580.

101. Wu X, Zhang Y, Ji M, Yang W, Deng T, Hou G, Shi L, Xun W. 2024. AhR Activation Ameliorates Intestinal Barrier Damage in Immunostressed Piglets by Regulating Intestinal Flora and Its Metabolism. Animals (Basel) 14.

102. Tonini M, Vicini R, Cervio E, De Ponti F, De Giorgio R, Barbara G, Stanghellini V, Dellabianca A, Sternini C. 2005. 5-HT7 receptors modulate peristalsis and accommodation in the guinea pig ileum. Gastroenterology 129:1557–66.

103. Wei L, Singh R, Ha SE, Martin AM, Jones LA, Jin B, Jorgensen BG, Zogg H, Chervo T, Gottfried-Blackmore A, Nguyen L, Habtezion A, Spencer NJ, Keating DJ, Sanders KM, Ro S. 2021. Serotonin Deficiency Is Associated With Delayed Gastric Emptying. Gastroenterology 160:2451–2466 e19.

104. Seo D, Patrick CJ, Kennealy PJ. 2008. Role of Serotonin and Dopamine System Interactions in the Neurobiology of Impulsive Aggression and its Comorbidity with other Clinical Disorders. Aggress Violent Behav 13:383–395.

105. Kwon YH, Wang H, Denou E, Ghia JE, Rossi L, Fontes ME, Bernier SP, Shajib MS, Banskota S, Collins SM, Surette MG, Khan WI. 2019. Modulation of Gut Microbiota Composition by Serotonin Signaling Influences Intestinal Immune Response and Susceptibility to Colitis. Cell Mol Gastroenterol Hepatol 7:709–728.

106. Liu Z, Ling Y, Peng Y, Han S, Ren Y, Jing Y, Fan W, Su Y, Mu C, Zhu W. 2023. Regulation of serotonin production by specific microbes from piglet gut. J Anim Sci Biotechnol 14:111.

107. Xie R, Jiang P, Lin L, Jiang J, Yu B, Rao J, Liu H, Wei W, Qiao Y. 2020. Oral treatment with Lactobacillus reuteri attenuates depressive-like behaviors and serotonin metabolism alterations induced by chronic social defeat stress. J Psychiatr Res 122:70–78.

108. Chong HX, Yusoff NAA, Hor YY, Lew LC, Jaafar MH, Choi SB, Yusoff MSB, Wahid N, Abdullah M, Zakaria N, Ong KL, Park YH, Liong MT. 2019. Lactobacillus plantarum DR7 alleviates stress and anxiety in adults: a randomised, double-blind, placebo-controlled study. Benef Microbes 10:355–373.

109. Vander Weele CM, Siciliano CA, Matthews GA, Namburi P, Izadmehr EM, Espinel IC, Nieh EH, Schut EHS, Padilla-Coreano N, Burgos-Robles A, Chang CJ, Kimchi EY, Beyeler A, Wichmann R, Wildes CP, Tye KM. 2018. Dopamine enhances signal-to-noise ratio in cortical-brainstem encoding of aversive stimuli. Nature 563:397–401.

